# Competition for food affects the strength of reproductive interference and its consequences for species coexistence

**DOI:** 10.1101/2023.11.09.566372

**Authors:** Miguel A. Cruz, Oscar Godoy, Inês Fragata, Vitor C. Sousa, Sara Magalhães, Flore Zélé

## Abstract

Competition for food and reproductive interference (negative interspecific sexual interactions) have been identified as major drivers of species exclusion. Still, how these biotic interactions jointly determine competitive dominance remains largely unknown. We tackle this by coupling population models and laboratory experiments with two sibling species of spider mites. Using experiments specifically designed to measure the single and combined effects of food competition and reproductive interference, we first show that the strength and symmetry of reproductive interference between species changes in presence of food competition. Next, we show that population models incorporating each type of interaction alone lead to markedly different predictions, from exclusion of the inferior competitor for food, to priority effects instead favouring the latter under the sole effect of reproductive interference. Moreover, accounting for the observed reduction in the strength of reproductive interference in the presence of food competition changes the threshold frequency determining the dominant competitor when both interactions are at play, from equal chances for the two species to exclude the other depending on their initial frequency, to favouring the superior competitor for food except when it is extremely rare. Finally, we showed that the model generates accurate predictions for population dynamics in an independent population cage experiment, indicating that our approach captures the most relevant processes governing the outcome of interactions between competing spider mite species. Altogether, our results suggest that trophic interactions can modulate sexual interactions, significantly impacting population dynamics and competitive outcomes. Hence, the joint consideration of food competition and reproductive interference is critical to accurately predict and understand species coexistence.

## 1 Introduction

How species that compete for common resources coexist is arguably one of the fundamental questions in ecology (Chesson, 2000; Huston, 1994; Hutchinson, 1961; Tilman, 1980). Modern coexistence theory posits that two competing species coexist when their niche differences, *i.e*. the degree to which intraspecific competition exceeds interspecific competition, are greater than their fitness differences, *i.e*., their differences in intrinsic growth rate weighted by their overall sensitivity to competition (Adler et al., 2007; Chesson, 2000, 2018; HilleRisLambers et al., 2012). A rich and vast literature also addresses how different abiotic and biotic factors affect the drivers, and ultimately, the outcomes of species competition and coexistence (Chesson, 1994; Dunson & Travis, 1991; McPeek, 2014; Spaak et al., 2021). In particular, competition for food often acts alongside other types of biotic interactions, but our knowledge of their joint effects is mostly limited to trophic ones, such as predation (Chesson & Kuang, 2008; Kotler & Holt, 1989; Shoemaker et al., 2020; Song et al., 2020) or parasitism (Hasik et al., 2023; Rovenolt & Tate, 2022; Terry et al., 2021), whereas the impact of other types of interactions, within the same trophic level, has received comparatively less attention to date.

Interspecific sexual interactions, owing to imperfect signal recognition, are relatively common between various assemblages of sexually reproducing species that generally also compete for food (Burns & Strauss, 2011; Kyogoku, 2015; Servedio & Hermisson, 2020; Webb et al., 2002; Weber & Strauss, 2016). Such interactions are commonly referred to as ‘reproductive interference’ as they generally have negative fitness consequences on at least one of the species involved (Gröning & Hochkirch, 2008). Because these interactions encompass a wide range of underlying mechanisms, varying from signal jamming, heterospecific rivalry, misdirected courtship and heterospecific copulations (incl. gamete wastage), up to the production of inviable or infertile hybrids (Gröning & Hochkirch, 2008), they include both direct (interference) and indirect (exploitative) competition for limited shared resources associated to species reproduction (*i.e*., breeding space, mates and/or gametes). Although historically reproductive interference has mostly been studied in speciation research, due to its central role in reproductive character displacement and reinforcement (Butlin, 1987; Servedio & Noor, 2003; Yamaguchi & Iwasa, 2015), it recently attracted growing interest in ecological research (Christie & Strauss, 2020; Cothran, 2015; Gómez-Llano et al., 2021, 2023; Grether et al., 2024; Weber & Strauss, 2016).

Theoretical studies posit that reproductive interference can be a major driver of competitive outcomes (Kishi & Nakazawa, 2013; Kuno, 1992; Schreiber et al., 2019; Yamamichi et al., 2023; Yoshimura & Clark, 1994). Because this interaction causes species to be more negatively impacted by heterospecifics than by conspecifics, it is predicted to promote positive frequency-dependence (*i.e*., contrarily to food competition it should disfavour the rarer species), and thus cannot act as a stabilizing mechanism (though see Gomez-Llano et al., 2018). This phenomenon, whereby the relative abundances of each species determine which one excludes the other and known as priority effects (Grainger et al., 2019), leads to reproductive interference driving exclusion faster than food competition (Kuno, 1992; Yoshimura & Clark, 1994). These effects can thus preclude long-term coexistence between ecologically equivalent species, as evidenced by an increasing number of laboratory and field studies (Hochkirch et al., 2007; Kishi, 2015; Kishi et al., 2009; Konishi & Takata, 2004; Liu et al., 2007; Ting & Cutter, 2018). Yet, as for other types of interactions (Amarasekare, 2002; Chesson, 2000), reproductive interference could promote coexistence via a trade-off with food competition if both interactions are asymmetric, such that the species most affected by food competition is less affected by reproductive interference or vice-versa (Kishi & Nakazawa, 2013; Schreiber et al., 2019). A limitation of prior work, however, is the assumption that food competition and reproductive interference operate independently from one another. In fact, several empirical studies revealed that, within species, food competition can affect the reproductive success of individuals (as better-fed males may be stronger competitors for mates, or even more attractive to females; Sigurjónsdóttir, 1984; Fisher & Rosenthal, 2006), and reciprocally, that different reproductive strategies may result in different levels of food competition (for instance, males that invest more in securing mates may be weaker competitors for food and vice-versa; Thiel & Dennenmoser, 2007). Such interplay between food competition and sexual interactions within species may also occur between species, such that food competition and reproductive interference might affect the strength of each other when simultaneously at play, as proposed by recent theory (Yamamichi et al., 2023).

In sum, both reproductive interference and competition for food can have strong impacts on species coexistence, but their joint effect remains largely experimentally unexplored. This knowledge gap may be due to the fact that, unlike other abiotic or biotic factors that can easily be singled out (*e.g*., temperature, salinity, competitors, parasites or predators), reproductive interference is part of the interaction between species that are simultaneously competing for food. Therefore, designing experiments that measure the impact of each of these interactions separately (which is needed as a control to evaluate their joint effect) is a true challenge. Although previous studies successfully uncoupled the effects of trophic and reproductive interactions on the population growth of co-occurring species (*Enallagma* damselflies, Gómez-Llano et al., 2023; and *Callosobruchus* beetles, Kawatsu & Kishi, 2018), and another assessed their joint effects on two parapatric *Streptanthus* jewelflower species (Christie & Strauss, 2020), none have, to our knowledge, directly compared the consequences of such interactions when acting separately *versus* jointly. Herbivorous spider mites represent an ideal system to do so, given their amenability to experimental manipulation of both competition for food (Fragata et al., 2022; Lu et al., 2018; Sarmento et al., 2011) and reproduction (incl. reproductive interference; Smith, 1975; Ben-David et al., 2009; Clemente et al., 2018; Sato & Alba, 2020).

Here, we studied the single and combined effects of reproductive interference and competition for food between two sibling species of spider mites, *Tetranychus urticae* and *T. cinnabarinus* (Kuang & Cheng, 1990; T. Li et al., 2009), also often referred to as the green and red forms of *T. urticae* (Auger et al., 2013). These two species have worldwide overlapping distributions and host plant ranges (Migeon & Dorkeld, 2023), sometimes being found on the same host plant (Lu et al., 2017; Zélé et al., 2018), on which they compete for food (Lu et al., 2017, 2018). These mite species also naturally engage in heterospecific sexual interactions (Smith, 1975), and suffer from strong reproductive interference due to variable reproductive incompatibilities and unfit hybridization (ranging from partial to complete hybrid sterility and hybrid breakdown; Dupont, 1979; de Boer, 1982; Sugasawa et al., 2002; Cruz et al., 2021; Xue et al., 2023). With this experimental system, we used recent modelling advances rooted in coexistence theory (Schreiber et al., 2019) and its recent extensions accounting for priority effects (Ke & Letten, 2018), and performed a set of experiments manipulating the presence or absence of food competition and reproductive interference. This combination of theory, modelling, and experiments allowed to estimate the strength of both types of interaction, when occurring alone or together, as well as to predict their single and joint contributions to competitive outcomes. Predicted dynamics of the two mite species interacting through both food competition and reproductive interference were then validated using data from an independent population experiment.

## 2 Materials and Methods

### 2.1 Biological materials

All populations used in this study were derived from a single source population of each of the two spider mite species, *Tetranychus urticae* (‘Tu’) and *T. cinnabarinus* (‘Tc’), collected in 2010 and 2013, respectively, from locations ca. 34 km apart (Cruz et al., 2021). Both source populations were free of any known endosymbiont and fully homozygous for the presence (Tu) or absence (Tc) of a recessive nonsynonymous mutation coding for pesticide resistance, which can be used to distinguish the individuals of each species in addition to stark differences in species-specific body colouration (Figure 1; see detailed explanations in Supplementary Materials Section S1). Due to incomplete pre-zygotic isolation (Cruz et al., 2025) and strong post-zygotic isolation (Cruz et al., 2021) between these two populations, strong reproductive interference is expected when they share the same environment (Smith, 1975). Indeed, females do not display any mate preference (non-assortative mating), whereas males of both species prefer to mate with Tc females (Cruz et al., 2025). Moreover, sex ratio distortion occurs in heterospecific crosses between Tu females and Tc males (with an overproduction of haploid sons instead of hybrid daughters, probably due to fertilization failure; Cruz et al., 2021), and both directions of heterospecific crosses result in fully sterile hybrids (most hybrid females do not lay eggs and the few eggs laid do not hatch; Cruz et al., 2021).

**Figure 1.**
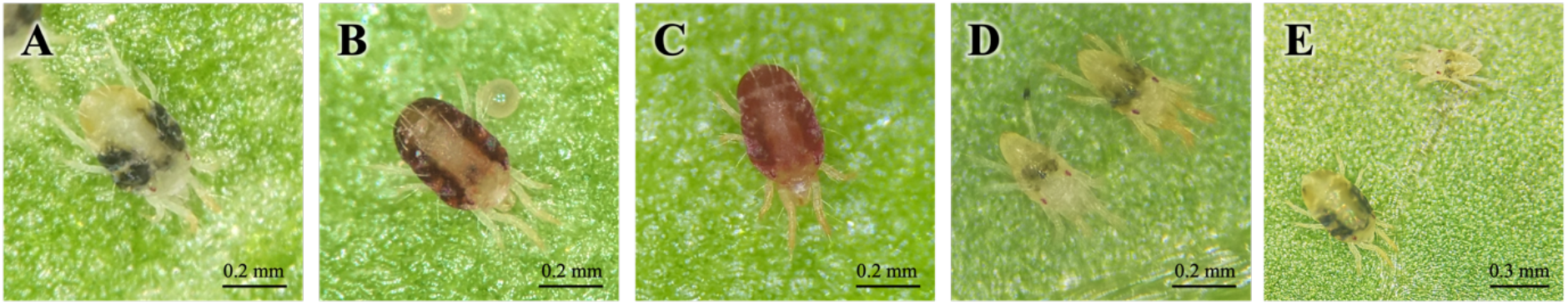
(A) *T. urticae*, (B) *T. cinnabarinus*, and (C) F1 hybrid females differ in their typical body colour, whereas (D) *T. urticae* (bottom left) and *T. cinnabarinus* (top right) males do not. (E) Adult female (bottom left) and male (top right) spider mites can be easily distinguished due to clear sexual dimorphism, including strong body size difference.

Prior to this study, five independent replicates were derived from each of the Tu and Tc source populations by transferring 200 adult females to a population cage containing two 14 days-old bean plants (*Phaseolus vulgaris*, cv. Contender seedlings obtained from Germisem, Oliveira do Hospital, Portugal). All replicate populations were then maintained under discrete generations at constant population size, by transferring 200 young adult females to new population cages every 14 days. Before performing each experiment, age cohorts were established from each of these replicate populations to obtain sufficient numbers of individuals of similar ages (see details in Section S2). Individuals from a given replicate population were always tested against those from the replicate population with the same label (*i.e*., Tu or Tc individuals of a given replicate were always tested against Tu or Tc individuals of the same corresponding replicate, and so on).

### 2.2 Measuring food competition, reproductive interference, and their combined effects

#### 2.2.1 Overview of the experimental procedures used to measure the separate and combined effects of each interaction

To uncouple the effects of food competition and reproductive interference on the growth of interacting species, as well as to measure the effect of reproductive interference in the presence of food competition, we performed a set of three different experiments, which are all fully detailed in Section S2. First, to quantify how food competition affected the per capita growth rate of both species in absence of reproductive interference (Experiment 1; Figure 2A), focal females that had already mated with conspecific males (hence were unexposed to reproductive interference) were exposed to a density gradient of conspecific and heterospecific competitors for food, and the number of their female offspring was counted to estimate their per capita growth rate (Hart et al., 2018). Second, to quantify the strength of reproductive interference while restricting competition for food (Experiment 2, Figure 2B), we counted the number of offspring produced by females isolated on individual food patches (hence unexposed to conspecific and/or heterospecific competitors for food) after they interacted sexually with both conspecific and heterospecific males. This was done over 2 generations to encompass the costs resulting from the production of sterile F1 hybrids. Third, to quantify the combined effect of food competition and reproductive interference on the growth rate of each species (Experiment 3; Figure 2C), we assessed the per capita production of daughters by females of each species when they interacted sexually and shared a common resource over 3 generations. In all three experiments, controls with no competitors for food nor heterospecific mates were also used to estimate the individual growth rate of each species in absence of interactions. Statistical analyses and offspring production data obtained in each experiment are reported in Section S3.

**Figure 2.**
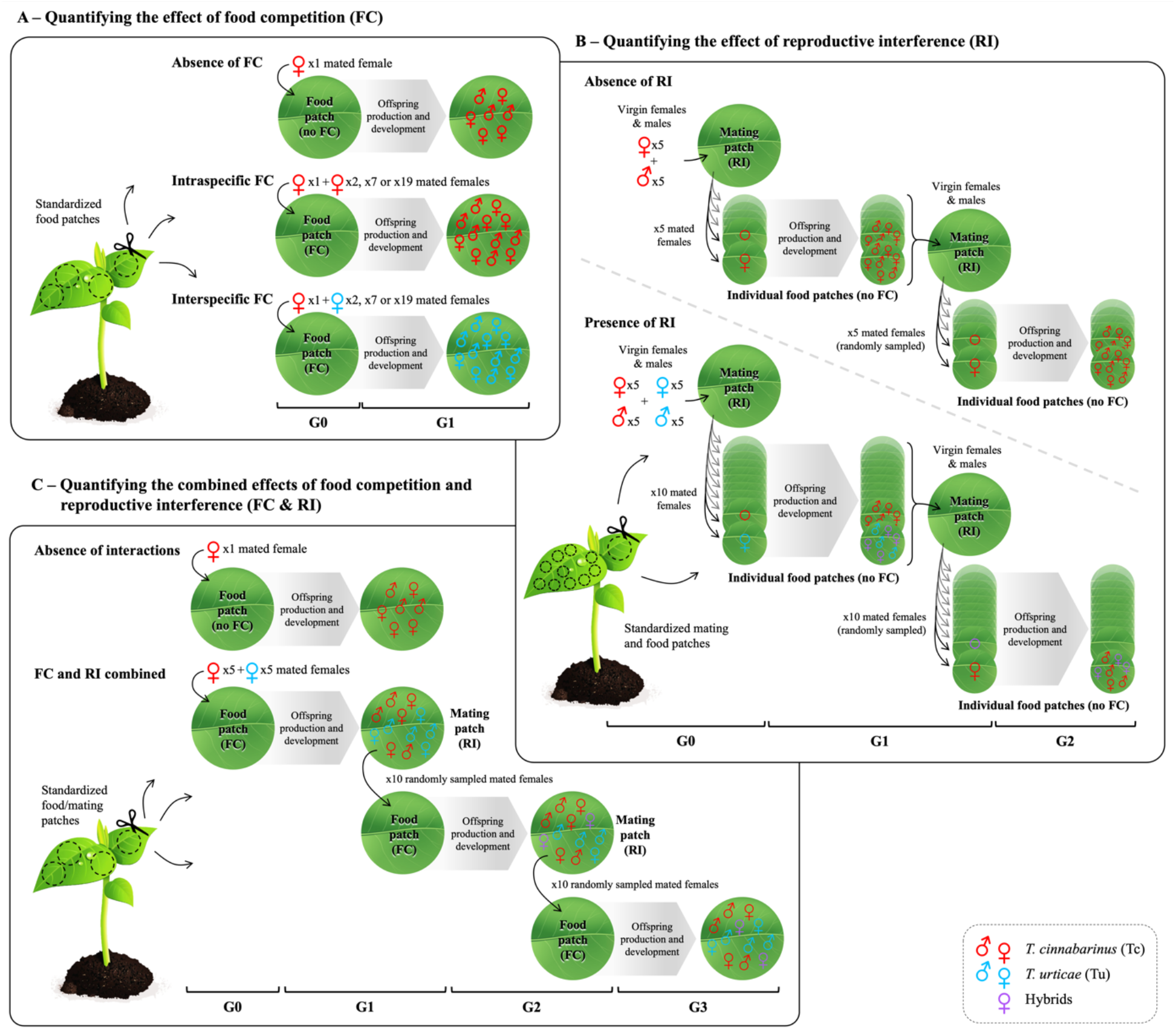
Experimental procedures used to estimate the individual and combined effects of food competition and reproductive interference. (A) Food competition was measured in absence of reproductive interference by placing adult females previously mated with conspecific males on food patches where they could lay eggs and their juveniles would develop. Intraspecific and interspecific competition are shown with Tc focal females as an example. (B) Reproductive interference was measured in absence of food competition by using common mating patches for virgin females and males of the two species, but individual food patches for ovipositing females and developing juveniles. The measures encompassed two consecutive generations (G0 to G2) to account for the costs resulting from F1 hybrid sterility. (C) Reproductive interference was measured in the presence of food competition across three consecutive generations (G1 to G3) by using common (food and mating) patches for the entire life cycle of the individuals. To correct for testing time, food competition in absence of reproductive interference was re-estimated during the first generation (G0 to G1), by using females mated with males from their own population at the onset of the experiment as in Experiment 1. The next generations of offspring (G2 to G3) allowed re-estimating reproductive interference in the presence of food competition, for two consecutive generations as in Experiment 2. FC: food competition; RI: reproductive interference; Red: *T. cinnabarinus* (Tc) females; Blue: *T. urticae* (Tu) females; Purple: hybrid females.

#### 2.2.2 Replication statement

The number of replicates performed in each of the different experiments in this study is provided in Table 1, and further detailed in Section S2. The scale of inference for all experiments is at the species level (though note that a single source population was used to establish the replicate populations of each species). The scales at which the factors of interest are applied are the leaf discs for the three first experiments, and the cage populations for the fourth experiment.

**Table 1.** Overview of replicates performed in all four experiments performed in this study.

| Experiment | Number of replicates at the appropriate scale |
| --- | --- |
| 1- Food competition only | 10 replicates<sup>1</sup> * 5 populations<sup>2</sup> * 2 focal species * 7 treatments = 700 total leaf discs |
| 2- Reproductive interference only | 10 replicates<sup>1</sup> * 5 populations<sup>2</sup> * 3 treatments (incl. 2 focal species) = 150 total leaf discs |
| 3- Food competition and reproductive interference | 10 replicates<sup>1</sup> * 5 populations<sup>2</sup> * 2 focal species for the “Interaction” treatment<br>30 replicates<sup>1</sup> * 5 populations<sup>2</sup> * 2 focal species for the “No interaction” treatment<br>= 100 + 300 total leaf discs |
| 4- Dynamics in cage populations | 5 cage populations<sup>2</sup> with the 2 species mixed (total population size of 400 within each cage) |

<sup>1</sup> experimental replicates
<sup>2</sup> replicate populations

### 2.3 Modelling approach to estimate interaction strengths from the experimental data and to predict the long-term outcome of species interactions

To estimate key interaction coefficients from data obtained in each experiment, we adapted a discrete-time population model developed by Schreiber et al. (2019), in which a reproductive interference component was added to the Beverton-Holt function used to predict population dynamics under food competition (Godoy & Levine, 2014). In this model, the number of individuals of a species *i* (*N*_*i*_) is predicted at each generation *t*, accounting for interactions with individuals of a species *j* (*N*_*j*_), with the following equation:

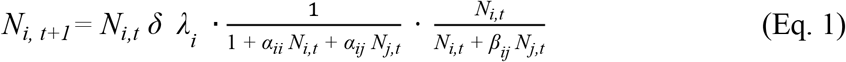

The first component of the expression (*i.e*., *N*_*i*,*t*_ *δ λ*_*i*_), refers to population growth in absence of limiting interactions for species *i*, with *δ* being the oviposition period in days and *λ*_*i*_ the per-capita daily intrinsic growth rate of species *i* in absence of any type of interaction. The middle component corresponds to the effect of food competition, with *α*_*ii*_ and *α*_*ij*_ being, respectively, the per capita effects of intraspecific and interspecific competition for food at generation *t*, hence with *N*_*i*,*t*_ and *N*_*j*,*t*_ competitors of species *i* and *j*, respectively. Finally, the third component of the expression represents the effect of reproductive interference, where *β*_*ij*_ is the per capita effect of interspecific sexual interactions with *N*_*j*,*t*_ individuals of species *j* on the growth rate of species *i*. We thus used a single parameter *β*_*ij*_ that encompasses the combined effects of all possible mechanisms underlying reproductive interference over multiple generations, instead of focusing on each of the different fitness components of reproductive interference possibly occurring at specific life stages of the organisms involved (see *e.g*., Gómez-Llano et al., 2023). This approach allows capturing the fitness costs resulting from various successive reproductive barriers being incomplete in a given system (*e.g*., heterospecific matings, fertilization failure and its consequences on offspring sex ratio, as well as hybrid sterility in the present system; Cruz et al., 2021, 2025) without being specific, as many other mechanisms may be involved (see Gröning & Hochkirch, 2008).

This population model was subsequently fit to the data obtained in each of the three experiments described above to estimate the values of the different parameter of the models in the different experimental conditions, as fully detailed in Section S4: the strength of food competition (*α*_*ii*_, *α*_*ij*_, *α*_*jj*_ and *α*_*ji*_) in absence of reproductive interference (*i.e*., when *β*_*ij*_ = *β*_*ji*_ = 0), the strength of reproductive interference (*β*_*ij*_ and *β*_*ji*_) in absence of food competition (*i.e*., when *α*_*ii*_ = *α*_*ij*_ = *α*_*jj*_ = *α*_*ji*_ = 0), and the strength of each interaction when they acted simultaneously (hence all alphas and betas).

Finally, estimated parameter values were used to predict the long-term outcome of trophic and/or sexual interactions between our populations, as fully described in Section S5. Briefly, because analytical solutions have not yet been derived for the full model (Schreiber et al., 2019), we focused instead on the three main conditions of interest for the present study:

- In the absence of reproductive interference (thus *β*_*ij*_ = *β*_*ji*_ = 0), the model simplifies to the Beverton-Holt function, and coexistence can be predicted using two key metrics (Chesson 2000): the average fitness differences, which refer to the competitive advantage of one species over the other; and the stabilizing niche differences, which represent the degree to which each species limits its own growth as compared to how it limits the growth of the competitor species (see Section S5).
- In the absence of food competition (thus *α*_*ii*_ = *α*_*ij*_ = *α*_*jj*_ = *α*_*ji*_ = 0), an analytical solution can be derived based on a third key metric: the relative strength of reproductive interference, which corresponds to the degree to which one species is more sensitive to reproductive interference than the other (as in Schreiber et al., 2019; see Section S5).
- When both interactions occur simultaneously, an analytical solution exists only for the specific case in which both species are equally affected by food competition, that is when this interaction is symmetrical (Schreiber et al., 2019). However, our results (see below) revealed that this is not the case in our system (as in many other systems such as annual plant communities; Allen-Perkins et al., 2023; Godoy & Levine, 2014). Hence, in this case, to determine the long-term consequences of the interaction strengths measured experimentally, we performed numerical analyses in which we varied the initial relative frequency and the total densities of individuals of each species (see detailed procedure in Section S5).

### 2.4 Empirical test of predicted population dynamics

To determine whether our estimated interaction coefficients enable accurate predictions of short-term population dynamics, we performed a larger-scale experiment (Experiment 4; Figure 3) in which we followed the relative frequencies of females of each species, as well as that of hybrid females, in cage populations over eight discrete generations, as fully detailed in Section S4. In addition, we adapted the previously used mathematical model (Eq. 1) to fit the procedure of the population cage experiment, that is, to model the random sampling of the females used to seed each new generation, as well as the asymmetrical production of sterile hybrid females due to fertilization failure occurring in our system (Cruz et al., 2021). This was done by implementing additional recursive steps in the model, and an additional parameter *θ*, corresponding to sex-ratio distortion in heterospecific crosses, as fully described in Section S6.

**Figure 3.**
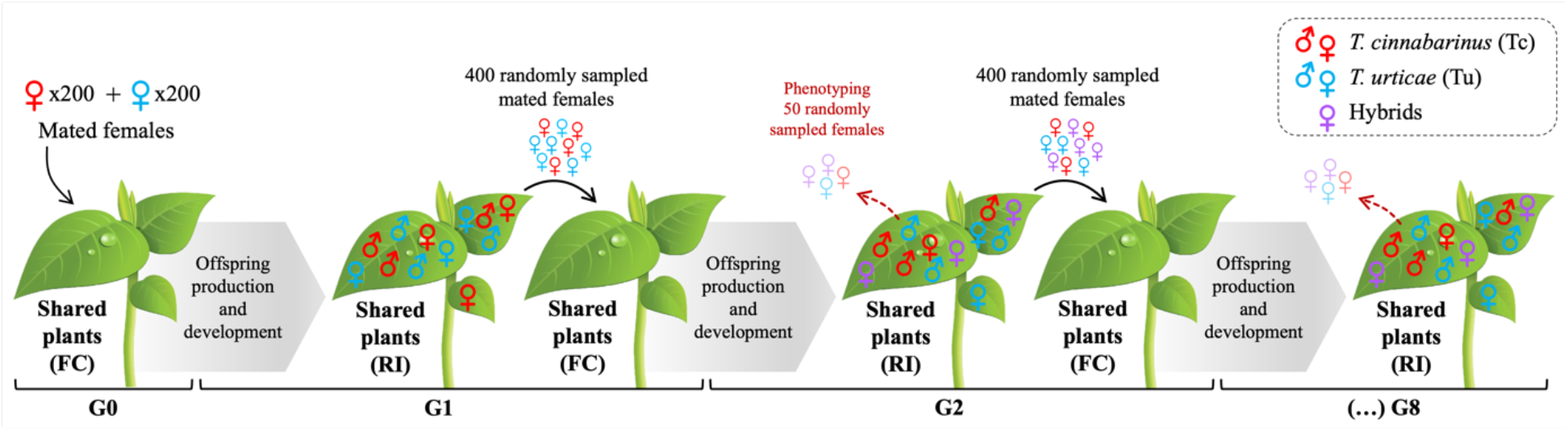
Experimental procedure used to measure population dynamics in cage populations over multiple generations. The same plants were used for the entire life cycle of individuals of both species. This procedure enabled food competition between females and developing juveniles, as well as reproductive interference between virgin adult individuals of both species. The figure depicts the initial installation of mated females (G0), the transfer of 400 randomly sampled mated females to start each subsequent discrete generation (G1 onward), as well as random sampling and phenotyping of 50 virgin females every two generations. FC: food competition; RI: reproductive interference.

Subsequently, we compared the observed proportion of each type of female with those predicted by the model, assuming either independent or combined effects of food competition and reproductive interference (by parameterising the model with estimates from Experiments 1 and 2, or from Experiment 3, respectively). Due to a discrepancy between the predicted and observed proportions of hybrid females (see Figure S4), we also made and compare additional predictions, assuming that hybrids were less affected by food competition than parental females by a range of scale factors of the *α* parameters in the equation used to predict hybrid production (see Section S6). Finally, we used linear regressions between the observed and predicted proportion of each types of females, for each of the different scenarios, to determine which model parameterisation leads to the most accurate predicted dynamics (see details in Section S6).

## 3 Results

### 3.1 Food competition affects the strength and symmetry of reproductive interference

Overall, all three experiments, in which females of each species were exposed to competition for food only (Experiment 1), reproductive interference only (Experiment 2), or both (Experiment 3), revealed subtle differences in population growth (Section S3), but large asymmetries in species intrinsic growth rates and interaction coefficients. First, intrinsic growth rates consistently differed between the two species in all 3 experiments: in the absence of competitors for food and heterospecific males, Tc females produced, on average, more offspring than Tu females (estimated coefficients *λ*_*Tc*_ > *λ*_*Tu*_ by *ca*. 25%; Figure 4A; Tables S1 and S2). Second, when species competed only for food (Experiment 1), both species were more sensitive to heterospecific than to conspecific competitors (estimated competition coefficients *α*_*TcTu*_ > *α*_*TcTc*_ by *ca*. 8%, and *α*_*TuTc*_ > *α*_*TuTu*_ by *ca*. 40%), but Tc females were generally more sensitive to competition than Tu females, regardless of competitor identity (*α*_*TcTc*_ > *α*_TuTu_ by *ca*. 140% and *α*_TcTu_ > *α*_*TuTc*_ by *ca*. 86%; Figure 4B; Table S1). Conversely, when the two species were only exposed to reproductive interference (Experiment 2), Tu females suffered more from the presence of heterospecifics than Tc females (estimated reproductive interference coefficients *β*_*TuTc*_ > *β*_*TcTu*_ by *ca*. 276%; Figure 4C; Table S1), suggesting that a trade-off between the two types of interaction could occur when both are at play.

**Figure 4.**
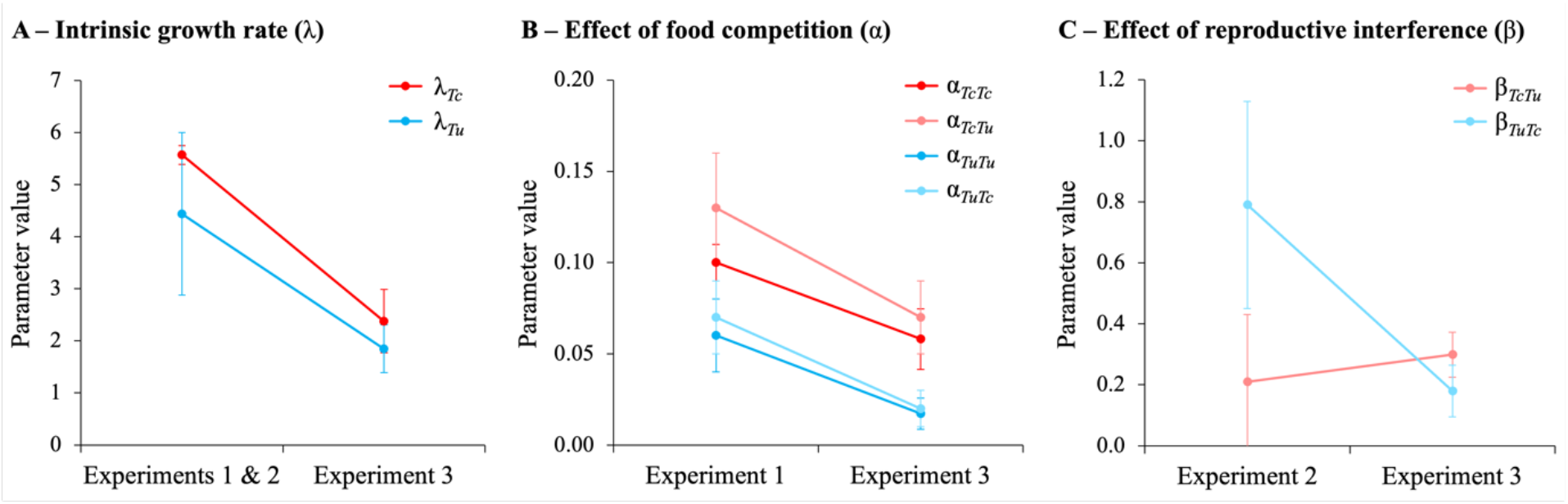
The strength and symmetry of reproductive interference change in the presence of food competition. (A) Intrinsic growth rate (*λ*) estimated as the mean number of daughters produced daily by single isolated females, averaged for Experiments 1 and 2, or measured in Experiment 3. (B) Per capita effect of food competition (*α*) estimated from Experiments 1 and 3. (C) Per capita effect of reproductive interference (*β*) estimated from Experiments 2 and 3. In all panels, dots show parameter values (± 95% confidence intervals) estimated across 5 replicate populations, dark and light colours represent within- and between-species effects, respectively, and parameter values for *T. cinnabarinus* (Tc) and *T. urticae* (Tu) are displayed in red and blue, respectively.

Finally, when both interactions occurred simultaneously (Experiment 3), we found lower intrinsic growth rates (*λ*_*Tc*_ and *λ*_*Tu*_) and strength of food competition (*α*_*TcTc*_, *α*_*TcTu*_, *α*_*TuTu*_, and *α*_*TuTc*_) than in the previous experiments, but both species were similarly affected by these changes: Tc females still produced more offspring (*λ*_*Tc*_ > *λ*_*Tu*_ by *ca*. 29%) and were more affected by food competition (*α*_*TcTc*_ > *α*_*TuTu*_ by *ca*. 200% and *α*_*TcTu*_ > *α*_*TuTc*_ by *ca*. 167%) than Tu females (Figure 4A, B; see also Table S2 *vs*. S1). However, we found a drastic change in the strength and symmetry of reproductive interference (*β*_*TcTu*_ and *β*_*TuTc*_): the sensitivity of Tc females to the presence of heterospecifics slightly increased and that of Tu females strongly decreased, such that Tu females switched from suffering more to suffering less from reproductive interference than Tc females (*β*_*TuTc*_ < *β*_*TcTu*_ by *ca*. 40%; Figure 4C). Hence, when simultaneously engaged in food competition and reproductive interference, Tu suffered less than Tc from both these interactions, and its population growth overall became higher than that of Tc from the second generation onwards (see Section S3).

### 3.2 The change in strength of reproductive interference due to food competition impacts theoretical predictions for coexistence

When we parameterised the population model with coefficients estimated from the small-scale experiment in which food competition acted alone (Experiment 1), we found negative niche differences in two replicate populations, as both species limited their competitor’s growth more than their own (*i.e*., each species was more affected by heterospecifics than by conspecifics). This should promote positive density-dependent effects (*i.e*., priority effects), whereby the species with higher relative abundance excludes the one with lower abundance. However, for three of the replicates and on average, we also observed large fitness differences between the two species, which rather lead to the prediction of Tu excluding Tc regardless of the relative abundance of the two species (Figure 5A).

**Figure 5.**
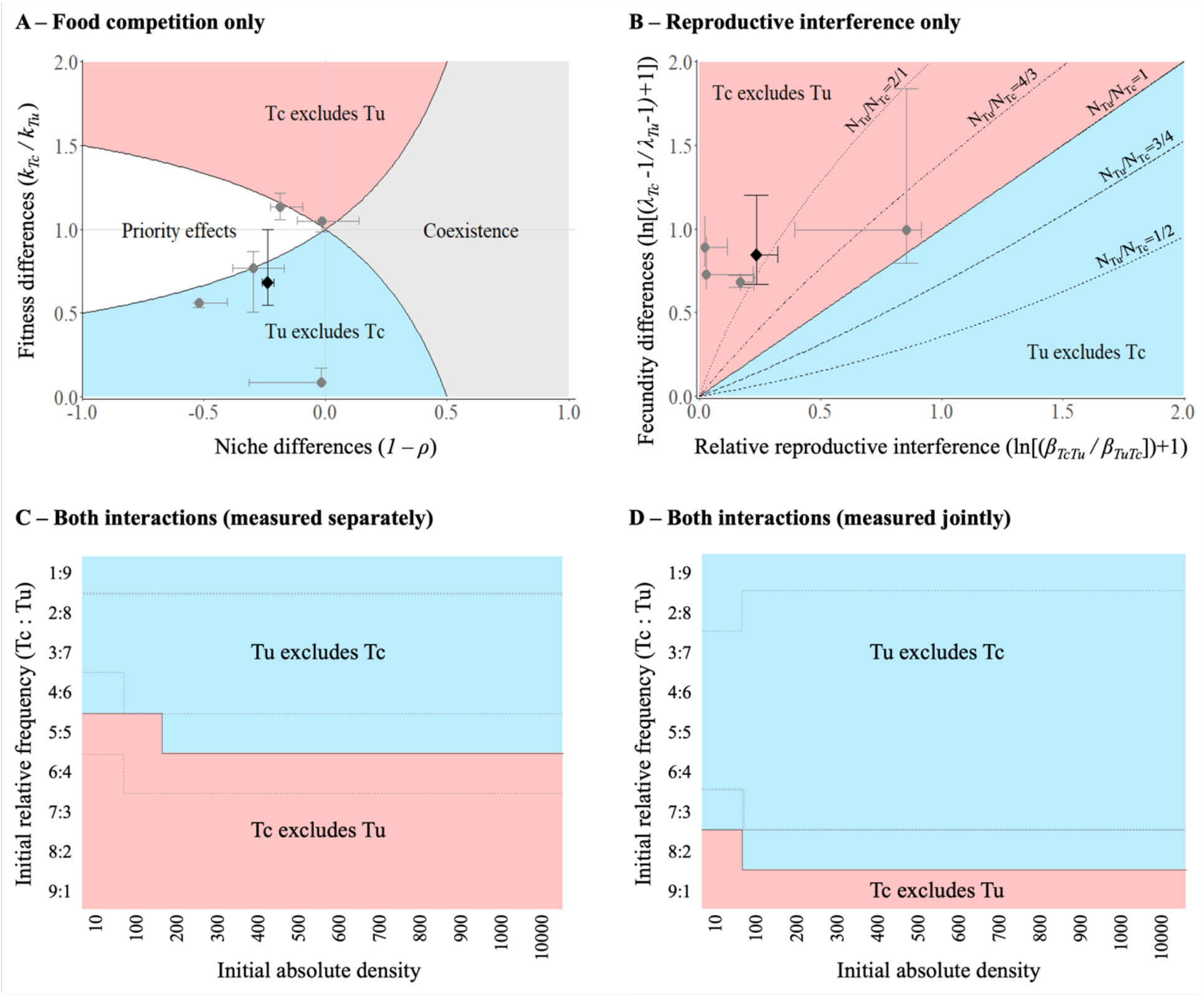
Changes in the strength of reproductive interference under food competition affect the predicted competitive outcomes. (A) Predictions depend on the niche and fitness differences between species when populations only compete for food. The black lines define the space in which species can coexist (grey), enter alternative stable states (priority effects; white), or where the species with higher fitness excludes the other (red or blue). (B) Predictions depend on the relative strength of reproductive interference and fecundity differences when populations only interact during reproduction. Coexistence is only possible if reproductive interference equalises the fecundity differences at equal initial frequency of both species (solid line: *N*_*Tu*_/*N*_*Tc*_ = 1), or if the initial frequency of the inferior species is sufficiently high to compensate for the combined effects of its fecundity disadvantage and/or higher sensitivity to reproductive interference (dotted lines from left to right: *N*_*Tu*_/*N*_*Tc*_ = 2/1, 4/3, 3/4, and 1/2). (C) and (D) Predictions after 20 simulated generations for different initial absolute densities and relative frequencies of the two species using interaction strengths measured when both types of interaction occurred separately or when they acted jointly, respectively. In A and B, grey circles and black diamonds are means (± 95% confidence intervals) for each replicate population and for all data, respectively. In C and D, dashed grey lines and solid black lines delimit the space in which each species excludes the other, using parameters estimated with each replicate population and with all data, respectively (but note that negative parameter values and confidence intervals are not displayed). In all panels, red and blue areas define the space in which *T. cinnabarinus* (Tc) excludes *T. urticae* (Tu), and vice-versa.

In absence of food competition, reproductive interference is expected to lead to priority effects (Schreiber et al., 2019), such that the outcome should depend not only on the combination of the relative strength of reproductive interference between the two species and their relative ‘fecundity difference’, but also on their initial relative frequency (*i.e*., the ratio between abundances of the two species). Indeed, different combinations of relative reproductive interference and fecundity differences determine different threshold frequencies above which one species is favoured over the other (Eq. S9; Figure 5B). In our system (*i.e*., with the parameter values we measured experimentally), we predicted that Tc should be favoured not only when both species are initially at equal relative frequency as in our experimental conditions, but also when Tc is less abundant than Tu (*i.e*., for all data and 3 replicate populations, it should be excluded only when its frequency drops below 33%; and below 50% for another replicate; Figure 5B). Thus, food competition and reproductive interference are predicted to lead to opposite outcomes when acting separately in our system.

Then, to determine how both interactions would jointly affect long-term population dynamics, we performed simulations with interaction strengths measured either when food competition and reproductive interference acted separately (using parameters estimated from Experiment 1 and 2) or jointly, hence when the strength and symmetry of reproductive interference changed in the presence of food competition (using parameters estimated from Experiment 3). In both cases, and similarly to our predictions for when reproductive interference acted alone, we found priority effects, with the identity of the species that excludes the other depending on initial conditions (both relative frequency and absolute density in this case; Figure 5C, D). However, accounting for changes in the strength and symmetry of reproductive interference under food competition drastically altered the threshold frequency determining which of the two species will be dominant (see Figure 5D *vs*. Figure 5C). When considering independent effects of the two interactions, we found that reproductive interference should counterbalance the asymmetries in food competition, with the weakest competitor for food (Tc) excluding the strongest competitor (Tu) when it is more abundant (threshold close to 50% on average, Figure 5C; see also Figures S1 and S2 for variation among replicate populations). In contrast, when the strength of reproductive interference is modulated by food competition, its buffering effect strongly decreases. Under this scenario, we found that Tu should exclude Tc except if its frequency drops below 20% on average (Figure 5D; see Figures S1 and S3 for variation among replicate populations).

### 3.3 The model accurately predicts dynamics in cage populations

To validate our hypothesis that accounting for changes in the strength of reproductive interference under food competition improves the accuracy of model predictions, we compared the dynamics observed in the population cage experiment (Experiment 4; Figure 6A) with those predicted by different fittings of the population model. In the experiment, all replicate populations showed consistent dynamics, with all Tc and hybrid females being excluded by the 6^th^ and 8^th^ generations, respectively (Figure 6B; Table S3). A similar pattern was also predicted by the model, both when accounting for independent effects of food competition and reproductive interference and when accounting for an asymmetrical reduction in reproductive interference under food competition. However, simulated population dynamics considering interaction strengths measured independently showed extremely large variance, indicating a strong uncertainty concerning which species should exclude the other, whereas predictions showed much lower variance when based on interaction strengths measured when food competition and reproductive interference acted jointly, with Tu always excluding Tc (*R*^*2*^ = 0.89; Figure S4). Thus, these results highlight the importance of incorporating the interplay between the effects of the two interactions to better predict population dynamics. Nevertheless, even when food competition and reproductive interference are acting simultaneously, the model predicted lower proportions of hybrids (from generations 2 to 6), and a faster exclusion of Tc as compared to experimental observations (Figure S4B *vs*. Figure 6B). Additional simulations in which we varied the sensitivity of hybrid females to food competition as compared to purebred females (Table S4) revealed that hybrid females might be at least 15 times less sensitive to food competition than purebred ones (Figure 6C) to obtain the best fit between observed and predicted dynamics (*R*^*2*^ = 0.97; Figure 6D) while only slightly increasing the AIC of the regression model (by 2.55; see Table S4).

**Figure 6.**
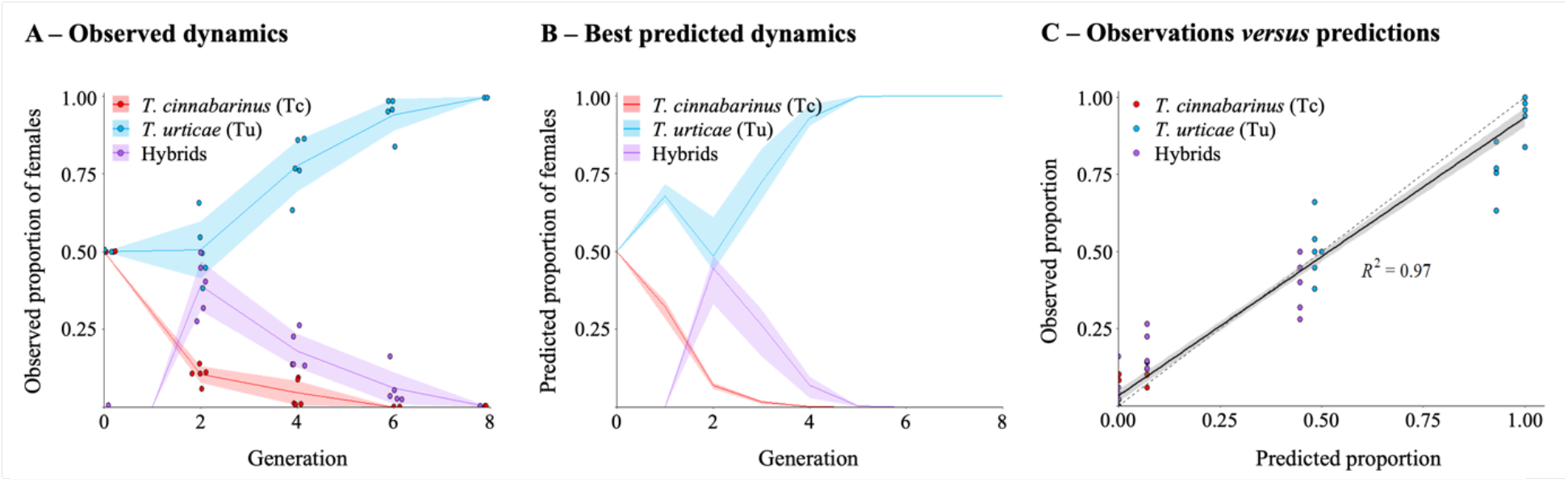
The model predictions accurately fit the observed dynamics. Proportions of *Tetranychus urticae* (Tu), *T. cinnabarinus* (Tc), and hybrid females (A) observed in experimental cage populations, over 8 generations, or (B) predicted assuming that food competition affects the strength of reproductive interference (parameters estimated from Experiment 3 across all replicate populations) and that hybrid females are 15 times less affected by food competition than purebred females. (C) Linear regression between observed and predicted proportions. In (A) and (C), dots show the observed proportions in each replicate population. In (A) and (B), coloured lines show the observed averages or predicted proportions across all replicate populations. In (C), the solid line shows the linear regression between observed and predicted proportions, and the dashed line shows the 1:1 relationship between values in the two axes. In all panels, shaded areas show 95% confidence intervals. Red: *T. cinnabarinus* (Tc) females; Blue: *T. urticae* (Tu) females; Purple: hybrid females.

## 4 Discussion

This study provides the first empirical evidence that the strength of reproductive interference can be affected by competition for food, such that the combined effects of the two types of interaction on population dynamics and competitive outcomes cannot be predicted by their independent action. Specifically, we show that, in absence of reproductive interference, *T. urticae* (Tu) is the stronger competitor for food and it is therefore predicted to invariably exclude *T. cinnabarinus* (Tc). Conversely, in absence of competition for food, Tu suffers more than Tc from reproductive interference and is thus predicted to be excluded, except if it is at least twice as abundant initially (due to positive-frequency dependence). Whereas a trade-off between competition for food and reproductive interference is, in the literature, predicted to enable stable coexistence (Kishi & Nakazawa, 2013; Schreiber et al., 2019), our simulations instead show that positive-frequency dependence driven by reproductive interference should remain when both types of interaction are simultaneously at play (the identity of the species that persists hinges upon their frequencies), be it with independent or interacting effects. When both types of interactions are assumed to have independent effects, food competition should balance out the advantage conferred to Tc by reproductive interference, with both species having the same likelihood to persist at an even initial frequency. However, when the strength of reproductive interference can be modulated by food competition, the superior competitor for food, Tu, becomes the least sensitive to reproductive interference. Our predictions accounting for such a change indicate the exclusion of Tc in all tested scenarios, except when it is initially extremely abundant relative to its competitor, Tu. The results obtained from an independent population cage experiment were largely compatible with this prediction, with Tc being systematically excluded when both species started at an even initial frequency. Our results therefore provide two straightforward lessons: first, food competition modulates reproductive interference, suggesting that these interactions are intertwined processes with non-independent effects on species coexistence, and second, modern coexistence theory (Schreiber et al., 2019; Yamamichi et al., 2023) is a suitable framework to predict their effects on the population dynamics and competitive outcomes of closely-related species.

Many studies have investigated the strength of reproductive isolation between Tu and Tc (Cruz et al., 2021, 2025; Murtaugh & Wrensch, 1978; Sugasawa et al., 2002; Xue et al., 2023). Still, reproductive interference and its consequences for population dynamics have barely been considered (but see Murtaugh and Wrensch, 1978). Here, in absence of competition for food, we found asymmetric reproductive interference between Tc and Tu, a result that is congruent with previous findings on pre- and post-mating reproductive isolation between the two populations used in this study (Cruz et al., 2021, 2025). Note, however, that the direction of the asymmetry in reproductive interference found here is unintuitive given the reproductive barriers identified in earlier studies on this system, in which crosses between Tu females and Tc males result in an overproduction of (Tu) male offspring due to fertilization failure, whereas males of both species preferentially mate with Tc females (Cruz et al., 2021, 2025). This should thus increase the risk of heterospecific mating for Tc females as compared to Tu ones, making them more likely to experience reproductive interference. The fact that we observed the opposite pattern here thus highlights that the relationship between reproductive barriers and reproductive interference is not as straightforward as one may think, and further studies are necessary to understand the complex interplay between the different mechanisms underlying reproductive interference (as in Gómez-Llano et al., 2023). Yet, the results of the present study are compatible with earlier studies using Chinese populations of the same species, both in the laboratory (Lu et al., 2017) and in the field (Lu et al., 2018). Indeed, these studies revealed that Tc consistently displaces Tu when at an even initial frequency, although this outcome is reversed in the presence of pesticides due to stronger pesticide resistance in Tu. In fact, any abiotic or biotic factor that affects the relative fitness of these two species, such as temperature (Gotoh et al., 2015; Lu et al., 2017; Riahi et al., 2013; Zou et al., 2018), host plants (Huo et al., 2021; Tomczyk et al., 1995; Witul & Kielkiewicz, 1999), or natural enemies (Takabayashi et al., 2000), should also play a role in determining their persistence in natural settings. Yet, no study so far aimed at disentangling the relative role of such factors in shaping competitive outcomes, let alone the role of different intrinsic factors.

Previous theoretical work predicted that food competition and reproductive interference acting in opposite directions (*i.e*., under a trade-off) may favour stable coexistence (Kishi & Nakazawa, 2013). Although we found that competition for food balances out the advantage conferred by reproductive interference to Tc in the current study, coexistence was not predicted in any scenario. Instead, simulated population dynamics revealed that positive frequency dependence driven by reproductive interference should occur even in the presence of food competition. This apparent discrepancy with previous predictions likely lies in the fact that we found negative niche differences in our system, a scenario yet unexplored despite several theoretical and empirical studies in the coexistence literature indicating that priority effects should be a common outcome of ecological interactions (Fragata et al., 2022; Ke & Letten, 2018; Song et al., 2021; Spaak et al., 2021). Negative niche differences in our study, however, could be a consequence of our experimental design, where only a single type of resource and no spatial heterogeneity was available, thereby severely precluding avoidance of competitors. Conversely, in natural populations, both species have similar but very vast host plant ranges (Migeon & Dorkeld, 2023), and food competition may shift these ranges in environments with more than a single plant species (Ferragut et al., 2013). Avoidance of competitors in natural populations could thus drive lower niche overlap between species, promoting coexistence at a broader scale (Wittmann & Fukami, 2018). Still, the evidence for spider mites avoiding interspecific competitors is mixed (Godinho et al., 2024; Pallini et al., 1997; Sato et al., 2016), hence it is not clear that they will be able to display this behaviour in all circumstances. Moreover, being crop pests, they are often exposed to a relatively homogeneous environment, in which the results outlined here are meaningful.

Our experiments were specifically designed to test for a change in the strength of reproductive interference in response to food competition. This possibility has not yet been investigated, neither theoretically nor empirically. There are, however, other forms by which these two interactions can affect each other. Indeed, recent theoretical work also suggests that changes in food competition intensity mediated by reproductive interference could drive coexistence by switching initially negative niche differences to positive (Yamamichi et al., 2023) even in the absence of alternative resources (*i.e*., due to behavioural and/or evolutionary changes; Kishi and Tsubaki 2014, Noriyuki and Osawa 2016, Ruokolainen and Hanski 2016). However, a thorough and systematic analysis of the joint effect of these two interactions is as yet in its infancy.

An experimental test of the combined effects of reproductive interference and competition for food revealed that these are not equivalent to the combination of their independent effects. Whereas the intrinsic growth rate of both species and their sensitivity to competition were overall consistently lower due to unknown environmental effects (Experiment 3), the intensity of reproductive interference became similar for both species in the presence of food competition, while it was stronger for Tu than for Tc in its absence (Experiment 2). This could be explained by a positive correlation between traits involved in each of these interactions, or by the same traits being involved in (and affected by) both interactions (Maan & Seehausen, 2011). For instance, body size is usually a key trait for both trophic and sexual interactions (Okuzaki et al., 2010). In spider mites, reduced food availability can negatively affect body size (G. Y. Li & Zhang, 2018), whereas larger males are generally superior competitors for mates (Enders, 1993; Potter et al., 1976), and larger females are preferred over smaller ones (Edward & Chapman, 2011; Zahradnik et al., 2008). Therefore, as Tu individuals are less impacted by food competition, they may retain larger body sizes, which should facilitate conspecific Tu matings (as Tu males may become more competitive) and thus reduce the strength of reproductive interference induced by Tc. Reduced food availability could also lead to slower offspring development in spider mites (Wilson, 1994), as in other arthropod systems (*e.g*., Teng & Apperson, 2000), such that Tu males, which are superior competitors for food, could develop faster and secure conspecific mates before Tc males become sexually mature, with similar consequences as just described above. Other mechanisms may also involve the production of signals of lower quality, or the production of fewer/smaller/less mobile gametes, etc. by the inferior competitor. Overall, food competition may thus affect trait values more severely in the offspring of the inferior competitor, which in turn become more affected by reproductive interference. Alternatively, a change in the strength of reproductive interference in the presence of food competition may simply arise via the effects that the latter has on population density, assuming that the strength of reproductive interference (β) varies with total population density (in the same way as the strength of food competition could change in the presence of reproductive interference if the α values are density-dependent, as proposed by Yamamichi et al.; 2023). However, for this hypothesis to explain our results, the β values would have to have a species-specific sensitivity to density.

Irrespective of the mechanism underlying the interplay between competitive and sexual interactions, our simulations revealed that changes in the strength of reproductive interference in response to food competition (as compared to its absence) lead to a strong increase in the threshold initial frequency under which the inferior competitor for food (Tc) cannot persist. Then, using independent data for model parameterisation and validation, we demonstrated the importance of accounting for this effect to accurately predict population dynamics, thereby further highlighting the importance of understanding how trophic and sexual interactions affect each other. This last piece of work also shows that a simple model capturing the demographic effects of reproductive interference with only two parameters (β and θ; *cf*. Methods), hence not explicitly modelling each of its underlying mechanisms (or ‘fitness components’; Kyogoku 2015), can generate very accurate predictions for the dynamics of two species in experimental cage populations. This ability to predict the system dynamics was further improved when accounting for the production of hybrid females that are less sensitive to food competition. While the ecological impact of hybrids is widely studied in the context of adaptation (Gow et al., 2007; Seehausen, 2004), the demographic impact of ‘unfit’ hybrids on parental species has been largely overlooked. In fact, an increased frequency of hybrids in parental populations (as predicted and observed here) could generate unpredicted changes in population dynamics, especially if they strongly compete for food and/or if they are highly attractive for males, regardless of their fertility. However, to our knowledge, it is yet unknown whether hybrid females in this system are recognized, or even preferred as potential mates by males of either species, or whether they are strong competitors for food. Nevertheless, consistent with observations made in our laboratory, our analyses revealed that they may require much less resources than purebred females, possibly because most of them do not produce eggs (Cruz et al., 2021).

Our study is a first but rigorous attempt to delve into the complexity of evaluating ecological interactions experimentally, while simultaneously accounting for competition for food and for mates. We have done the most we could perform given our logistical limitations, but larger studies could address additional aspects not covered in our study to fully comprehend their complex interactions. For instance, we could design experiments to determine whether the relationship between each species’ sensitivity to intraspecific and interspecific competition also changes in the presence of reproductive interference, or whether and how reproductive interference affects the intensity of food competition (*i.e*., the opposite of what we tested here) as proposed by Yamamichi et al. (2023). For instance, within-species male mating harassment (Oku, 2009) could either decrease or increase the intensity of food competition, respectively by reducing female feeding time as they invest more time in evading male mating attempts (Bancroft & Margolies, 1996), or by increasing spatial aggregation of females in order to create refuges from males (Yamamichi et al., 2023). Furthermore, the model could be developed to account for within-species negative reproductive interactions, such as male harassment, which may also play an important role in determining the strength of reproductive interference between populations (Kyogoku & Sota, 2017). Interactions between conspecific males and females may even be altered by the presence of heterospecifics, potentially resulting in facilitation rather than interference (Gomez-Llano et al., 2018). Such a theoretical development could ultimately allow for an expansion of the concept of niche differences not only to trophic, but also to sexual, interactions, leading to an even more powerful integration into a general coexistence framework (Gómez-Llano et al., 2021).

In conclusion, our study reveals that food competition can affect the strength of reproductive interference, with significant consequences for the dynamics of competing species. Given that reproductive interference is expected to often occur between species that compete for food (Servedio & Hermisson, 2020), such outcome is likely a general feature of many ecological systems involving taxa that incidentally engage in sexual interactions (Weber & Strauss, 2016), and it may have major consequences for the study of ecological coexistence (Germain et al., 2018; Gómez-Llano et al., 2021). Indeed, our study shows that addressing the effect of each interaction type alone might be insufficient to accurately predict population dynamics. This is because the co-occurrence of both types of interactions can change the shape of the trade-off between reproductive interference and competition for food, unbalancing the priority of one species over another in systems with negative niche differences (while potentially hampering coexistence in systems with positive niche differences). Understanding this interplay between feeding and sexual interactions is also crucial for speciation research, as the likelihood and duration of coexistence between closely-related species will determine the opportunities for reinforcement of pre-zygotic reproductive isolation. Our findings may thus have far-reaching consequences for a recently growing field at the interface between speciation and coexistence theory (Boussens-Dumon & Llaurens, 2021; Germain et al., 2018; Grether et al., 2020; Kyogoku & Kokko, 2020; Kyogoku & Wheatcroft, 2020). Finally, despite species interactions being intensively studied for more than a century as a major determinant of species distribution and competitive outcomes, our results collectively show that we are still in the infancy of understanding how different mechanisms interact to determine the population dynamics of interacting species. Combining theoretical and empirical approaches is key to unveil how different types of interactions jointly impact species coexistence or exclusion.

## Acknowledgements

We are grateful to Inês Santos, Cátia Eira and Miguel Cunha for the maintenance of the spider mite populations, to Lucie de Sousa for growing the plants, and to André Mira and Alexandra Balola for assistance with data collection. We are also grateful to Emanuel A. Fronhofer and Jhelam N. Deshpande for helpful discussions on the modelling work, and to Patrice David and Enric Frago for their advises and comments on the manuscript. This work was funded by an ERC Consolidator Grant (COMPCON, GA 725419) attributed to SM. MC was funded through an FCT PhD fellowship (SFRH/BD/136454/2018) and IF through a Junior researcher contract (CEECIND/02616/2018). OG acknowledges financial support provided by the Spanish Ministry of Economy and Competitiveness and by the European Social Fund through the project UCCO (CNS2023-144337). VCS acknowledges financial support provided by FCT (CEECINST/00032/2018/CP1523/CT0008), including national funds to cE3c (UIDB/00329/2021). This is contribution ISEM-2024-XXX of the Institute of Evolutionary Science of Montpellier (ISEM). For the purpose of Open Access, a CC-BY 4.0 public copyright licence has been applied by the authors to the present document and will be applied to all subsequent versions up to the Author Accepted Manuscript arising from this submission.

## Open Research statement

All datasets and R scripts used in this study are available at Zenodo (https://doi.org/10.5281/zenodo.10086125).

## Author Contributions

MC, VS, OG, SM and FZ designed the experiments. MC collected data. MC and FZ performed modelling work and analysed data with advises and scripts from IF and OG. MC and FZ wrote the first draft of the manuscript, and all authors contributed substantially to revisions.

## Conflict of interest

The authors declare that they have no conflict of interest with the content of this article.

## Notes

https://doi.org/10.5281/zenodo.10086125

### Competing Interest Statement

The authors have declared no competing interest.

### Summary of Updates

Figures have been updated to more clearly separate methodology from results, and several details in the procedure of the experiments and parameter estimation have been moved to supplementary materials to trim manuscript length. Additionally, sections of the Introduction and Discussion have been expanded and/or clarified.

