## Supplementary Materials for "Competition for food affects the strength of reproductive interference and its consequences for species coexistence"

**Section S1. Procedures used to obtain the *Wolbachia*-free and genetically distinguishable source populations used in the current study.**

The two source populations used in this study were initially naturally infected with *Wolbachia* (though uninfected with any other known endosymbiont of spider mites). Therefore, we first treated them with antibiotics to obtain *Wolbachia*-free populations (Cruz et al. 2021) in order to ensure that reproductive interference in our experiments would result only from host-intrinsic factors. To achieve this, 400 females of each population laid eggs on bean leaves placed on cotton soaked with a 0.05% rifampicin solution. From these eggs, 265 *T. urticae* (Tu) and 308 *T. cinnabarinus* (Tc) adult females were obtained and confirmed to be uninfected through multiplex PCR (Zélé et al. 2018). These individuals were used as the founders for the source populations.

Additionally, to enable unambiguous genetic identification of mites from the two source populations, we used a recessive, nonsynonymous, mutation (I1017F) in the chitin synthase 1 (CHS1) gene, known to confer resistance to the pesticide etoxazole (Van Leeuwen et al. 2012). This resistance mutation was shown to not carry fitness costs in our standard experimental conditions (Costa et al. 2024), which made it a practical molecular marker (Rodrigues et al. 2020, Costa et al. 2023). The Tu population was initially heterozygous for this gene, whereas the Tc population lacked this mutation. Thus, to ensure that the Tu population did not contain any susceptible individuals prior to performing experiments, 1650 females were allowed to lay eggs on bean leaves placed on cotton soaked with a ~0.5% etoxazole solution (concentration above LC100; Van Leeuwen et al. 2012). This resulted in 377 homozygous pesticide-resistant daughters, which were propagated to build the fully etoxazole-resistant Tu source population used in this study.

**Section S2. Detailed procedures used to experimentally measure food competition, reproductive interference, and their combined effects.**

*Age cohorts prior to performing experiments*

To obtain a sufficiently large number of individuals of roughly the same age for each experiment, all replicate populations of the two species were expanded ca. 4 weeks before performing an experiment, by transferring 100 adult females to a separate population cage containing one fresh bean plant. Eleven to 15 days later, age cohorts were created from each of these additional boxes. Groups of 75 (for Experiment 1, food competition alone; see below) or 50 (for Experiment 2, reproductive interference alone, and Experiment 3, food competition and reproductive interference; see below) mated females, as well as 50 virgin females (to produce only males for Experiment 2), were collected from each replicate population and installed to lay eggs for 3 days on detached bean leaves placed on water-soaked cotton in Petri dishes. For Experiments 1 and 3, adult mated female offspring were collected from each age cohort of each replicate population 14 days later, and directly used on the first day of the experiments. For Experiment 2, virgin female offspring were collected 11-12 days later (two days prior to the start of the experiment), while they were still deutonymphs undergoing their last moulting stage. They were placed for two days in groups of 30 without males on *ca.* 9 $cm^2$ bean leaf fragments, such that they emerged as adults while remaining virgin. Virgin males were directly collected from the cohorts on the first day of the experiment. Population maintenance and all experiments were performed under the same controlled laboratory conditions (24±2ºC, 70% humidity, 16/8h L/D).

*Experiment 1: Food competition in absence of reproductive interference*

To experimentally establish a density gradient to measure food competition (Hart et al., 2018), one adult mated female of the Tu and/or Tc population was installed as a focal individual on a *ca.* 9 $cm^2$ bean leaf disc placed on water-soaked cotton contained in a Petri dish to lay eggs for 3 days, either alone, or together with 2, 7, or 19 conspecific or heterospecific female competitors (hence seven different 'treatments' were performed; Figure 2A in the main article file). At the end of the egg-laying period, all females were discarded, and 11 days later (14 days after the experiment onset) the number of adult daughters that were produced by the focal females and had survived competition on each leaf disc was counted (in the case of intraspecific competition, the total number of daughters per disc was divided by the initial female density, whereas for interspecific competition, offspring of the focal female were identified based on body colouration; see Figure 1 in the main article file). We focus on the production of daughters as we assume that the contribution of spider mite males to food competition is negligible (Bonte & Bafort, 2019; Fragata et al., 2022; see also the size difference between adult males and females in Figure 1E in the main article file).

*Experiment 2: Reproductive interference in absence of food competition*

We installed 5 virgin females and 5 virgin males from each cohort of each replicate population on a *ca.* 9 $cm^2$ bean leaf disc placed on water-soaked cotton contained in a Petri dish (*i.e.*, a 'big' mating patch to prevent individuals from incidentally leaving the patch and drowning), either in the absence of heterospecifics ('control treatments'; one for each species), or with 5 virgin females and 5 virgin males from a cohort of the other species, to allow for reproductive interference between the two (*i.e.*, 'reproductive interference treatment'; Figure 2B in the main article file). Individuals could freely mate for 2 days before males were discarded and each female was isolated on 2.5 $cm^2$ bean leaf discs (individual patches) to lay eggs for 1 day. The offspring produced by each female thus developed in the absence of food competition with the offspring of other females. Nine days later, when all female offspring were at (or about to enter) their last moulting stage and most male offspring were already adults, all offspring within a given experimental replicate were merged on a fresh mating patch, where they could emerge as adults and interact (incl. mating) for 3 days. Then, the same number of females as initially used in each treatment (10 for the reproductive interference treatment and 5 for the control treatments) were randomly sampled in each mating patch and isolated on individual patches to lay eggs for 1 day. After 11 days, the number of adult daughters in each individual patch was registered and the total number of daughters of each species per replicate was computed (*i.e.*, species identification was based on body colouration in this experiment; see Figure 1 in the main article file). Offspring production was registered over two generations to encompass all costs resulting from reproductive interference, including offspring sex ratio distortion and the production of sterile F1 hybrids.

*Experiment 3: Combined effects of food competition and reproductive interference*
We installed 5 Tc and 5 Tu adult mated females on a *ca.* 9 $cm^2$ bean leaf disc to lay eggs for 3 days (*i.e.*, 'interactions' discs; Figure 2C in the main article file), at which point these adult females were discarded. Eleven days later (14 days after the experiment onset), the number of adult daughters of each species was counted and up to 10 adult females were randomly selected and transferred to a fresh bean leaf disc (*ca.* 9 $cm^2$) to start the next generation. Female offspring was counted 11 days later, as before, and the entire procedure was repeated until the third generation. Because all females used to initiate the experiment were already mated to conspecific males, the first generation was only affected by food competition and no hybrids were formed. Then, because the females that were randomly sampled at the first and second generations were already mated to either conspecific, heterospecific, or both types of males, the second and third generations were affected by both food competition and reproductive interference, and hybrid daughters were also produced (and counted). Finally, females of either species were also installed individually on *ca.* 9 $cm^2$ bean leaf discs to estimate their individual growth rate in this experiment (*i.e.*, 'absence of interactions' discs, one for each species). These females could lay eggs for 3 days and their daughters were counted 11 days later. As in the previous experiments, the identification of females of each species (as well as their hybrids) was based on body colouration (Figure 1A-C in the main article file).

*Experiment 4: Dynamics in cage populations*
In order to determine the efficiency of the model in making accurate predictions based on previously estimated parameters (see modelling methods in Section S6), we followed the dynamics of replicated populations in experimental cages across 8 generations. Prior to the experiment, the Tu and Tc source populations were expanded by placing 3 to 4 mite-infested bean plants in a large population cage (28 x 39 x 28 cm) with 4 fresh bean plants. The experiment started 14 days later by installing 200 mated Tu females with 200 mated Tc females in 5 independent population cages (14 x 20 x 14 cm), each containing 4 fresh bean plants (*ca.* 624 $cm^2$ total leaf area). This amount of food was chosen because it should trigger moderate food competition for a total of 400 females (*i.e.*, it is equivalent to a density of *ca.* 6 individuals on a *ca.* 9 $cm^2$ bean leaf disc in Experiment 1; see Results). Nine days later, 2 fresh bean plants were added in each of the cages to avoid complete resource depletion, and 5 more days were given to the offspring of both species to become adult and interact sexually. Fourteen days after the onset of the experiment, 200 mated females were randomly sampled from each of the two newest plants (hence a total of 400 females) of each cage, and transferred to a new population cage to start the next generation. This entire procedure was repeated at each generation for each of the 5 independent replicates (*i.e.*, cage populations), until complete exclusion of one of the species and of the hybrids. The proportions of individuals of each species, as well as their hybrids, were estimated every 2 generations through phenotyping of a subset of female offspring immediately before fresh plants were added on day 9. For this purpose, 80 females undergoing their deutonymph stage were randomly sampled in the experimental cage and installed on a *ca.* 9 $cm^2$ bean leaf fragment to emerge as virgin adult females. After 3 days, 50 of them were installed on individual 2.5 $cm^2$ leaf discs placed on cotton soaked with ~0.5% etoxazole solution. Females were given three days to lay eggs before being discarded, and five days later, each disc was checked for eggs and juveniles. As ingestion of etoxazole by susceptible adult females prevents their eggs from hatching (Van Leeuwen et al., 2012), pesticide application in our system was used to identify adult females according to the following criteria: if at least one egg hatched, the female was identified as resistant to etoxazole, hence as a Tu female; if no egg hatched, the female was identified as susceptible to etoxazole, hence as a Tc female. Finally, because *ca.* 99% of hybrid females in this system are sterile (Cruz et al., 2021), non-ovipositing females were identified as hybrids. This procedure was used to account for the fact that a hypothetical, yet unlikely, very low level of gene flow could hamper the efficiency of phenotyping based on body colouration over multiple generations (conversely to the Experiments 2 and 3, which were performed up to the production of hypothetical F2 hybrids only).

*Detailed description of the replications performed in each experiment*
The Experiment 1 (food competition only) was performed in 10 independent temporal blocks, installed over the course of 2 weeks (5 blocks per week, installed on 5 consecutive days). Each competition treatment (focal female installed either alone, or together with 2, 7, or 19 conspecific or heterospecific female competitors) and replicate population were tested simultaneously, with one experimental replicate (*i.e.*, leaf disc) of each being carried out per block. Hence, 10 experimental replicates (one per block) were performed per replicate population of each species and per treatment, for a total of 700 replicates (10 experimental replicates*5 replicate populations*2 species*7 treatments) obtained at the end of the experiment.

The Experiment 2 (reproductive interference only) was performed in 5 independent temporal blocks installed over the course of 5 weeks, with a maximum of 2 blocks on consecutive days. Two experimental replicates (*i.e.*, mating patch) per treatment (Tc alone, Tu alone, or both species together) and replicate population were carried out simultaneously within each block. Hence, 10 experimental replicates (2 experimental replicates*5 blocks) were performed per replicate population and per treatment, for a total of 150 experimental replicates (10 experimental replicates*5 replicate populations*3 treatments) obtained at the end of the experiment.

The Experiment 3 (food competition and reproductive interference) was carried out in 3 independent temporal blocks, installed over the course of 2 weeks (2 blocks installed on consecutive days at the start of the first week and the third block installed at the start of the second week). For the 'interactions' treatment (*i.e.*, containing both species together), 4 experimental replicates of each of the five replicate populations were performed in the first block and 3 were carried out in the remaining two blocks, for a total of 50 discs (10 experimental replicates*5 replicate populations). The 'Absence of interactions' treatment (*i.e.*, containing single females of either species) was performed concurrently with the first and third blocks of the experiment, in 15 experimental replicates per block, for a total of 150 discs (30 experimental replicates*5 replicate populations).

The Experiment 4 (dynamics in cage populations) was carried out in a single experimental block, containing 5 independent experimental replicates (*i.e.*, boxes). These consisted of cage populations with a mix of both species at a total population size of 400 adult females.

**Section S3. Methods and results of statistical analyses performed on the data obtained in each of the three experiments testing for the combined or separate effects of food competition and reproductive interference on offspring production.**

*Procedure used for statistical analyses*

To statistically investigate the overall combined or separate effects of food competition and reproductive interference on offspring production in Experiments 1 to 3, the daily per capita number of daughters obtained for each focal species at the last generation of each experiment (generation 1 for Experiment 1, 2 for Experiment 2, and 3 for Experiment 3) was analysed using generalized linear mixed models (glmmTMB package; Brooks et al. 2017) with a Poisson or with a negative binomial error distribution and/or accounting for zero inflation when overdispersion was detected ('family' argument chosen based on AIC-based model comparisons; see selected/used models in Table I). For all analyses, maximal models containing the complete set of explanatory variables (see Table I) were subsequently simplified by sequentially eliminating non-significant terms to establish a minimal model (Crawley 2007). The significance of explanatory variables was established using chi-square tests with the *anova* function. Significant values given in the text are for the minimal model, whereas non-significant values correspond to those obtained before deletion of the variable of interest from the model (Crawley 2007).

**Table I. Description of the statistical models used in the analyses of offspring production in the three experiments.** For all analyses, the variable of interest is the per capita daily daughter production, either alone or in the presence of heterospecifics. This variable corresponds to the number of daughters divided by the number of initial females and by the number of days of oviposition. For each analysed dataset (or data subset), the "N" column gives number of observations in each analysis. "Maximal model" provides the complete set of fixed explanatory variables initially included in the model, from which non-significant fixed factors were sequentially removed to establish a minimal model. Experimental block and replicate population were fit as random explanatory variables in all models. focal: species of the focal individuals; competition: type of competition assessed (intraspecific or interspecific); heterospecifics: whether heterospecifics were present or not. The "Family" column indicates the error distribution used for each analysis, with p: Poisson (variance = $\mu$), nb1: negative binomial with a linear parametrization (variance = $\mu + \mu*\exp(\eta)$), and nb2: negative binomial with a quadratic parametrization (variance of $\mu + \mu^2/\exp(\eta)$), where $\mu$ is the predicted mean and $\eta$ is the linear predictor from the dispersion model; 'zi' prefix indicates that the model accounts for zero inflated data distribution).

| Model no. and corresponding data subset | N | Maximal model | Minimal model | Family |
|---|---|---|---|---|
| **Experiment 1** | | | | |
| 1.1 All data | 699 | focal * competition * density | focal * competition * density | zip |
| 1.2 Females exposed to conspecifics | 400 | focal * density | focal * density | zip |
| 1.3 Females exposed to heterospecifics | 299 | focal * density | focal * density | p |
| **Experiment 2** | | | | |
| 2.1 All data | 200 | focal * heterospecifics | focal * heterospecifics | zinb1 |
| 2.2 Females not exposed to heterospecifics | 100 | focal | 1 | zip |
| 2.3 Females exposed to heterospecifics | 100 | focal | focal | nb1 |
| **Experiment 3** | | | | |
| 3.1 All data | 100 | focal | 1 | nb2 |

*Results*

In the first experiment, we found that differences between the two focal species depended on the type of competition they were exposed to ($\chi^2_1 = 4.82$, $P = 0.03$, model 1.1 in Table I, Figure I.A): overall Tc females produced less offspring than Tu females across all competitor densities, but differences between the two species were smaller when these were exposed to intraspecific (*ca.* 1.5% difference; $\chi^2_1 = 4.29$, $P = 0.04$; model 1.2) than to interspecific competitors (*ca.* 4% difference; $\chi^2_1 = 25.3$, $P < 0.0001$; model 1.3).

In the second experiment, we found that differences in offspring production by the two species varied in the presence or absence of heterospecifics (interaction focal * heterospecifics: $\chi^2_1 = 14.99$, $P = 0.0001$; model 2.1, Figure I.B). Indeed, when heterospecifics were absent, both species produced similar numbers of offspring ($\chi^2_1 = 1.19$, $P = 0.28$; model 2.2), but Tc produced *ca.* 107% more offspring than Tu when either species was exposed to heterospecifics ($\chi^2_1 = 21.21$, $P < 0.0001$; model 2.3).

Finally, statistical analyses revealed no significant differences in offspring production between the two species at the third generation of Experiment 3 ($\chi^2_1 = 0.84$, $P = 0.36$; model 3.1, Figure I.C).

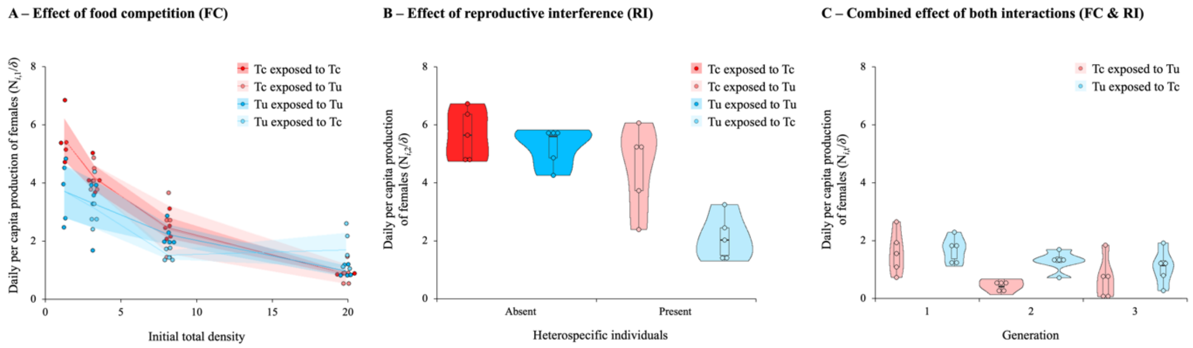

**Figure I. The effects of food competition, reproductive interference, and a combination of both, on offspring production.** (A) Number of females produced daily under increasing densities of either conspecific or heterospecific competitors (dark and light colours, respectively) in Experiment 1. Individual points show the mean number of females produced daily by a single female (per capita) for each of 5 different replicate populations, whereas lines show the average across all replicates and shaded areas represent the 95% confidence intervals. (B) Number of females produced daily at the second generation of Experiment 2 (G2), when focal species grew in the absence or presence of heterospecific individuals (dark and light colours, respectively) on a common mating patch, but in absence of food competition. Individual points show the mean number of females produced daily by a single female (per capita) for each of 5 different replicate populations, whereas violin plots show the mean distribution across all replicates, and boxplots within each violin plot show the median, 1st and 3rd quartiles, and whisker length corresponds to 1.5 times the interquartile range. (C) Number of females produced daily at the third generation of Experiment 3 (G3), in the absence or presence of heterospecific individuals (dark and light colours, respectively), when individuals shared the same food and mating patches over 3 generations. Individual points show the mean number of females produced daily by a single female (per capita) for each of 5 different replicate populations, whereas violin plots show the mean distribution across all replicates, and boxplots within each violin plot show the median, 1st and 3rd quartiles, and whisker length corresponds to 1.5 times the interquartile range. FC: food competition; RI: reproductive interference; Red: *T. cinnabarinus* (Tc) females; Blue: *T. urticae* (Tu) females; Purple: hybrid females.

**Section S4. Detailed procedure used to estimate the parameters of the mathematical model.**

All parameter estimations were done in the R statistical software (v4.2.2) using custom scripts available at Zenodo (https://doi.org/10.5281/zenodo.10086125). Values of $\lambda$ were computed as the mean (and 95% confidence interval) daily production of daughters ($\lambda_i = N_{i,\, t+1}/\delta$) in the control treatments of each experiment (single females in Experiments 1 and 3, or conspecific females in Experiment 2). For subsequent analyses, parameter estimations and predictions, we either used the average of the $\lambda$ estimates obtained from Experiments 1 and 2 (in which each type of interaction was measured independently), or used directly the $\lambda$ estimated from Experiment 3 (in which both types of interactions were measured simultaneously). Values of the $\alpha$ and $\beta$ interaction coefficients were estimated using a maximum likelihood approach similar to that of Godoy and Levine (2014). Briefly, the method consisted in fitting the log-transformed number of daughters observed experimentally into Eq. 1, along with fixed values of $\lambda$ obtained from the experiments. Note that, because our estimation of $\lambda$ already included the effects that a focal individual has on itself, we excluded the focal individual in the food competition component of Eq. 1 for the estimations of the $\alpha$ and $\beta$ coefficients (*i.e.*, we considered only the number of neighbour females). Model fitting then allowed identifying the parameter values (randomly sampled within the normal density function *dnorm*) and associated standard deviations (from which the 95% confidence intervals were obtained) that enable the best fit between the model predictions and the observed data (*i.e.*, by minimizing the negative log-likelihoods; function *optim*). This was done with a simulated Nelder-Mead algorithm (Nelder & Mead, 1965), instead of the 'L-BFGS-B' method used by Godoy and Levine (2014), to avoid false convergence issues. For all $\alpha$ and $\beta$ estimations, we used an unrealistic starting value of 4, then repeated the estimations with a more realistic value of 0.1 to verify the consistency in the outcomes.

When only competition for food could occur (Experiment 1), the value of $\beta$ was set to 0 when fitting Eq. 1, which is equivalent to entirely removing the reproductive interference component from the model, simplifying it to the classical Beverton-Holt function. Then, both intraspecific ($\alpha_{ii}$ and $\alpha_{jj}$) and interspecific ($\alpha_{ij}$ and $\alpha_{ji}$) competition coefficients were estimated using only data from the 'intraspecific' and 'interspecific' competition treatments, respectively (*i.e.*, offspring produced by focal females that had already mated with conspecific males and were exposed to an increasing gradient of conspecific or heterospecific females in the neighbourhood).

When only reproductive interference occurred (Experiment 2), the values of all $\alpha$ parameters were set to 0 when fitting Eq. 1, which is equivalent to removing the food competition component from the model. Then, the reproductive interference coefficients $\beta_{ij}$ and $\beta_{ji}$ were estimated simultaneously using the data obtained after two generations in the treatment in which 5 virgin females and 5 virgin males of both species were initially mixed on a same mating patch, but females were subsequently isolated on individual food patches after they interacted sexually.

Finally, when both types of interactions occurred (Experiment 3), we could not estimate both $\alpha$ and $\beta$ values simultaneously due to practical limitations, hence we used a nested approach instead to avoid structural non-identifiability (Raue et al., 2009) as well as overfitting (as in Matías et al., 2018). Competition coefficients ($\alpha$) were first estimated using only data obtained from the first generation (during which reproductive interference did not occur), whereas reproductive interference coefficients ($\beta$) were estimated using the data obtained from the generations 2 and 3 (hence over 2 generations, exactly as for $\beta_{ij}$ and $\beta_{ji}$ estimations in Experiment 2). In this case, the value of the previously estimated $\alpha$'s was fixed in the model. A logistic limitation with this approach is that, by design, we only assessed changes in the $\beta$ values (reproductive interference) due to the occurrence of food competition (by comparing the values obtained from Experiments 2 and 3), but not changes in the $\alpha$ values (food competition) due to the occurrence of reproductive interference. Also, because in Experiment 3 we could not independently measure the effect of intra- and interspecific competition, as we did in Experiment 1 (*i.e.*, we could not test different absolute densities of intra- and interspecific competitors due to the complexity of the experimental design when subsequently allowing for reproductive interference over

2 additional generations), it was not possible to estimate all $\alpha$ values simultaneously. To mathematically circumvent this issue, we forced the $\alpha_{ii}/\alpha_{ij}$ and $\alpha_{jj}/\alpha_{ji}$ ratios to be equal to the $\alpha_{ii}/\alpha_{ij}$ and $\alpha_{jj}/\alpha_{ji}$ ratios obtained in Experiment 1, respectively. We followed this procedure assuming that, for each species (for each replicate population, as well as for all data independently of the replicate), the relationship between values of intra- and interspecific coefficients would be similar between Experiment 1 and 3.

**Section S5. Theoretical approach to predict the long-term outcomes of species interactions.**

*Outcomes of food competition in the absence of reproductive interference*

When only food competition occurs (*i.e.*, reproductive interference is absent, hence $\beta_{ij} = \beta_{ji} = 0$), the model simplifies to the Beverton-Holt function, for which analytical solutions can be easily computed using two key metrics: the average fitness difference and the niche overlap between the two species (Chesson, 2000).

The average fitness difference between two species refers to the competitive advantage of one species over the other. It corresponds to the product of two ratios: the 'demographic ratio' that describes the ratio of intrinsic growth rates between species in the absence of competitors, and a 'competitive response ratio' that describes the degree to which one species is more resistant to competition than the other (Chesson, 2012). It is thus computed using the estimated $\lambda$ and $\alpha$ parameters as follows:

$$\frac{k_j}{k_i} = \frac{\lambda_j - 1}{\lambda_i - 1} \cdot \frac{\sqrt{\alpha_{ij}\alpha_{ii}}}{\sqrt{\alpha_{ji}\alpha_{jj}}} \qquad \text{(Eq. S1)}$$

When the value of $k_j/k_i$ is above 1, species $j$ is superior (*i.e.*, it should exclude species $i$ in the absence of reproductive interference and under complete niche overlap; see below); when it is below 1, species $i$ is superior; and when it is close to 1, the two species are equivalent competitors. Thus, a reduction in fitness difference has an equalising effect on the population dynamics of interacting species (Chesson, 2000).

The niche overlap between species, $\rho$, represents the degree to which each population limits its own growth as compared to how it limits the growth of the competitor species (Chesson, 2012). It is computed using the estimated $\alpha$ parameters as follows:

$$\rho = \sqrt{\frac{\alpha_{ij}\alpha_{ji}}{\alpha_{jj}\alpha_{ii}}} \qquad \text{(Eq. S2)}$$

When the value of $\rho$ is below 1, a species limits its own growth more than that of its competitor, favouring coexistence. However, when the value approaches 1 (full niche overlap), species limit each other's growth as much as they limit their own, and coexistence is hampered (one of the species will always exclude the other). Thus, niche difference, expressed as $1 - \rho$, has a stabilising effect on population interactions (Chesson, 2000). Finally, if niche overlap is greater than one, population dynamics may be dominated by priority effects, such that the order of arrival or initial frequencies of individuals from the two species become the determinants for which one will exclude the other (Ke & Letten, 2018). Note that the competition coefficients in Eq. S2 (the $\alpha$ parameters) are phenomenological, but we assume in our work that species limit each other mainly through food competition.

Theory predicts that, for species to coexist, they must either be equivalent competitors, or niche differences must be high enough as to overcome fitness differences (Chesson, 2012), hence that:

$$\rho < \frac{k_j}{k_i} < \frac{1}{\rho} \qquad \text{(Eq. S3)}$$

*Outcomes of reproductive interference in the absence of food competition*

When only reproductive interference occurs (*i.e.*, food competition is absent, hence $\alpha_{ii} = \alpha_{jj} = \alpha_{ij} = \alpha_{ji} = 0$), the outcome can be computed as a function of another key metric previously described by Schreiber et al. (2019).

The relative strength of reproductive interference, $\gamma$, represents the degree to which one species is more sensitive to reproductive interference than the other. It is computed using the estimated $\beta$ parameters as follows:

$$\gamma = \frac{\beta_{ji}}{\beta_{ij}} \quad \text{(Eq. S4)}$$

When the value of $\gamma$ is above 1, species $j$ is superior (*i.e.*, it should exclude species $i$ in absence of food competition, provided that both populations have equal intrinsic growth rate; see below); when $\gamma$ is below 1, species $i$ is superior; and when it is close to 1, the two populations are equally (symmetrically) affected by reproductive interference. Thus, as for fitness differences, symmetrical impacts of relative reproductive interference have an equalising effect.

Using this metric, the analytical solution to predict the long-term outcome of reproductive interference when it occurs in the absence of competition for food can be derived as follows:

In absence of food competition, the growth rate of each species is given by (*i.e.*, Eq. 1 simplifies as):

$$\frac{N_{i,\,t+1}}{N_{i,\,t}} = \frac{(\lambda_i \delta)\, N_{i,\,t}}{N_{i,\,t} + \beta_{ij} N_{j,\,t}} \quad \text{(Eq. S5)}$$

Thus, at equilibrium (*i.e.*, when $N_{i,\,t+1} = N_{i,\,t}$), the strength of reproductive interference experienced by species $i$ will depend on the relative frequency of each species:

$$\beta_{ij} = \frac{N_i}{N_j} \cdot \left(\lambda_j\, \delta - 1\right) \quad \text{(Eq. S6)}$$

And the growth rate of both species will reach an equilibrium when:

$$\frac{N_i}{N_j} = \frac{\beta_{ij}}{(\lambda_i\, \delta - 1)} = \frac{(\lambda_j\, \delta - 1)}{\beta_{ji}} \quad \text{(Eq. S7)}$$

Which allows computing the relative growth of each species as a function of the relative strength of reproductive interference:

$$\left(\frac{N_i}{N_j}\right)^2 = \frac{(\lambda_j\, \delta - 1)}{(\lambda_i\, \delta - 1)} \cdot \frac{1}{\gamma} \quad \text{(Eq. S8)}$$

Hence, coexistence is possible (though unstable) when:

$$\frac{N_i}{N_j} = \sqrt{\frac{(\lambda_j\, \delta) - 1}{(\lambda_i\, \delta) - 1} \cdot \frac{1}{\gamma}} \quad \text{(Eq. S9)}$$

Coexistence is thus frequency-dependent (Schreiber et al., 2019): it only occurs when the relative frequency of the two species balances the ratio between their 'fecundity' differences (*i.e.*, fitness differences in absence of competition) and relative reproductive interference. In the specific case where both species have the same initial relative density ($N_i = N_j$), the species with the lower fecundity must be proportionally less affected by reproductive interference. If these conditions are not satisfied, alternative stable states can be found (*i.e.*, the system is driven by priority effects), with the species that displays a fitness advantage and/or lower sensitivity to reproductive interference excluding the other, even when outnumbered by its competitor to some extent (*i.e.*, species $i$ excludes species $j$ when $N_j(\lambda_j\text{-}1)/\gamma\, N_i(\lambda_i\text{-}1) < 1$).

*Outcomes of food competition and reproductive interference when both interactions are at play*

When two species simultaneously engage in food competition and reproductive interference, analytical solutions are only available for the specific case in which the species are ecologically equivalent ($\alpha_{ii} = \alpha_{jj}$ and $\alpha_{ij} = \alpha_{ji}$; Schreiber et al., 2019). Because this assumption is hard to meet in nature and indeed it was not observed in our system (see Results, Figure 4B), and it is beyond the scope of the present study to derive analytical solutions for the general case, we opted for numerical analyses. Briefly, we parameterised the population model (Eq. 1 in the main article file) with the parameter values estimated when both types of interaction acted independently (Experiments 1 and 2) or simultaneously (Experiment 3), along with different sets of mock parameter values that are predicted to allow for coexistence (as controls) when the two species only engage in food competition or in reproductive interference, or simultaneously engage in both interactions (see Figure II). We simulated population dynamics for a wide range of initial absolute densities (total population size ranging from 100 to 1000 individuals in increments of 100, as well as 10 and 10000 individuals) and relative frequencies (ranging from 1:9 to 9:1 individuals of each species) of individuals of the two species. Simulations were run over 20 generations, which was the minimum number of generations necessary for the system to reach stability in all studied scenarios.

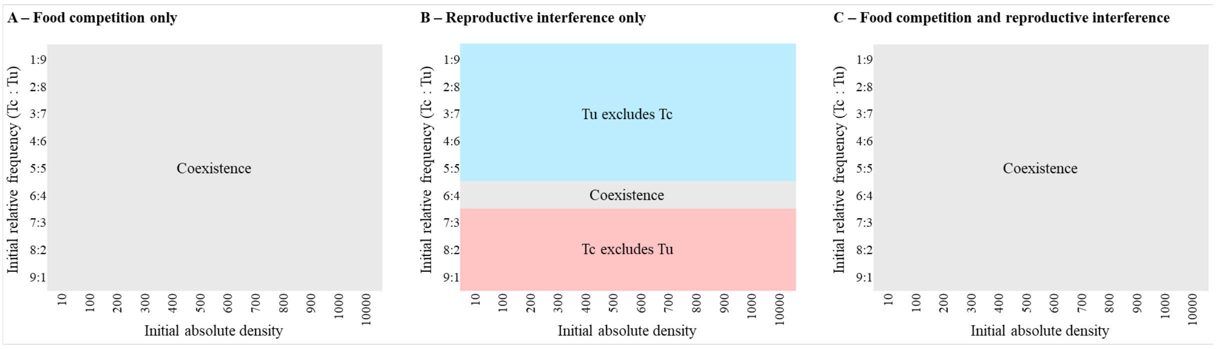

**Figure II. Summary outcome of population dynamics simulated with mock parameter values as a control for the simulation method.** In all panels, the heat map shows the species that is predicted to remain after 300 generations, for different initial absolute densities and relative frequencies of both species. This was done for different sets of mock parameter values that should allow (A) stable coexistence when species only engage in food competition ($\lambda_{Tu} = 4$; $\lambda_{Tc} = 2$; $\alpha_{TuTu} = 0.5$; $\alpha_{TcTc} = 0.3$; $\alpha_{TuTc} = 0.2$; $\alpha_{TcTu} = 0.1$; $\beta_{TuTc} = \beta_{TcTu} = 0$), (B) unstable coexistence when species only engage in reproductive interference ($\lambda_{Tu} = 4$; $\lambda_{Tc} = 2$; $\alpha_{TuTu} = \alpha_{TcTc} = \alpha_{TuTc} = \alpha_{TcTu} = 0$; $\beta_{TuTc} = 2$; $\beta_{TcTu} = 1.5$), and (C) stable coexistence when species simultaneously engaged in both interactions, due to a trade-off between both interactions in a system with positive niche differences ($\lambda_{Tu} = 100$; $\lambda_{Tc} = 160$; $\alpha_{TuTu} = \alpha_{TcTc} = 1$; $\alpha_{TuTc} = \alpha_{TcTu} = 0.1$; $\beta_{TuTc} = 0.1$; $\beta_{TcTu} = 0.3$). Red: only Tc remains, Tu is excluded; Blue: only Tu remains, Tc is excluded; Grey: both species stably coexist.

**Section S6. Detailed procedure used to predict species dynamics in cage populations, for model parameterisation and for comparison of predicted *vs.* observed dynamics.**

*Adjustment of the mathematical model to fit the procedure used in the 'cage population' experiment (Experiment 4; see Section S2).*

To predict the proportion of Tu, Tc, and hybrid females in cage populations for multiple generations, we adapted and extended the previously described model (Eq. 1 in the main article file) as follows:

First, the number of purebred female offspring of species $i$ ($N_i$) produced by a given number of sampled females $i$ ($S_i$), and introduced in a new experimental cage at each generation, is given by:

$$N_{i,\,t+1} = S_{i,t}\lambda_i\delta \cdot \frac{1}{1+\alpha_{ii}S_{i,t}+\alpha_{ij}S_{j,t}} \cdot \frac{S_{i,t}}{S_{i,t}+\beta_{ij}S_{j,t}} \quad \text{(Eq. S10)}$$

Note that the only difference between this equation and Eq. 1 is that $N_{i,\,t+1}$ is computed as a function of previously sampled females ($S_i$ and $S_j$), instead of previously produced females ($N_i$ and $N_j$).

Second, the number of hybrid female offspring ($N_h$) produced at each generation by the sampled purebred females of both species ($S_i$ and $S_j$) as:

$$N_{h,\,t+1} = \frac{S_{i,t}\lambda_i\delta}{1+\alpha_{ii}S_{i,t}+\alpha_{ij}S_{j,t}} \cdot \theta_{ij} \cdot \left(1-\frac{S_{i,t}}{S_{i,t}+\beta_{ij}S_{j,t}}\right) + \frac{S_{j,t}\lambda_j\delta}{1+\alpha_{jj}S_{j,t}+\alpha_{ji}S_{i,t}} \cdot \theta_{ji} \cdot \left(1-\frac{S_{j,t}}{S_{j,t}+\beta_{ji}S_{i,t}}\right) \quad \text{(Eq. S11)}$$

In this equation, the number of hybrid females produced by each species of sampled purebred females is given by the left and right terms of the expression, respectively. In both cases, this number corresponds to the number of female offspring becoming hybrids instead of purebred females due to reproductive interference. Because, in Eqs. 1 and S10, the 'reproductive interference factor' represents the proportion of female offspring 'escaping' reproductive interference (*i.e.*, successfully developing as purebred female), the opposite of this proportion represents the proportion of female offspring 'affected' by reproductive interference, hence (potentially) developing as hybrid females. However, not all female offspring produced by heterospecifically-mated mothers become hybrids in our system, as some can develop as males instead (likely due to fertilization failure; Cruz et al. 2021). Thus, to account for sex-ratio distortion in heterospecific crosses, we used an additional parameter $\theta$, which corresponds to the proportion of hybrid females produced by a purebred female $i$ mated with a male $j$ ($\theta_{ij}$), or by a purebred female $j$ mated with a male $i$ ($\theta_{ji}$). Finally, note that Eq. S10 and Eq. S11 assume that food competition only occurs among purebred females, not with hybrid females. Indeed, due to near-complete hybrid sterility, these latter consume very few resources as they do not expend energy producing eggs (personal observation). Moreover, the model assumes that purebred females producing hybrids are equally affected by food competition as those whose offspring escape reproductive interference (*i.e.*, the food competition factors are identical in both equations; but see different tested scenarios below).

Finally, the proportion of Tu, Tc, and hybrid female offspring ($P_i$, $P_j$, and $P_h$, respectively) in the population at each generation was computed as:

$$P_{i,\,t} = \frac{N_{i,t}}{N_{i,t}+N_{j,t}+N_{h,t}} \quad \text{(Eq. S12)}$$

And the number of Tu, Tc, or hybrid females randomly sampled ($S_i$, $S_j$, and $S_h$, respectively) in the population to start the next generation was computed as:

$$S_{i,\,t} = P_{i,t} \cdot S_{tot} \quad \text{(Eq. S13)}$$

Where $S_{tot}$ corresponds to the total number of female offspring sampled in each experimental cage to start the next generation ($S_{tot} = 400$ in our experiment).

*Parameterisation of the model and comparison of predicted vs. observed dynamics*

To determine the mechanisms that could best explain the observed dynamics in our population cage experiment, we parameterised the model, in which food competition and reproductive interference are always simultaneously at play, following different scenarios. The first two scenarios aimed at determining whether using parameter values estimated when food competition and reproductive interference acted separately (Experiments 1 and 2) *vs.* simultaneously (Experiment 3) significantly affected the ability of the model to accurately predict population dynamics. Then, building upon the second scenario (estimates from Experiment 3), we considered a third scenario, where hybrid females were less sensitive to food competition than purebred females. The model so far assumed the same levels of food competition between the offspring of purebred females that suffered from, or escaped, reproductive interference. However, this assumption may be unrealistic if hybrid females require fewer resources than purebred females during their development. Note that although not evaluated, we repeatedly observed very low levels of leaf damage in patches hosting developing and adult hybrid females relative to those hosting only purebred females, even at similar female densities. Thus, to determine whether a lower sensitivity of hybrid females to food competition would improve the predictions of the model, we divided all $\alpha$'s in Eq. S10 by a scale factor, testing every integer ranging from 2 (*i.e.*, hybrids suffering half as much from food competition as purebred females) to 30. To determine which scale factor of $\alpha$ enables the best model predictions as compared to the observed data, we used a binomial GLM (generalized linear model with a binomial error structure; package *lme4* in R), in which we fitted the observed proportions of individuals of each species and of the hybrids as the response variable, and the predicted proportions of offspring as fixed explanatory variable. We then compared the fit of each set of predictions based on the R-squared values, intercepts and regression coefficients of the linear regressions. For all tested scenarios, the model was adapted to account for the fact that no reproductive interference occurred, thus no hybrids were produced, during the first generation of the experiment. Values taken for $\theta_{ij}$ and $\theta_{ji}$ were the inverse of the $MD_{corr}$ indexes in Cruz et al (2021), which are estimates of the proportion of offspring developing as a haploid male instead of a female in interspecific crosses between our source populations. Upper and lower confidence intervals for the observed proportions of Tu, Tc, and hybrid females were calculated from the data, whereas those for the predicted proportions were calculated as the upper and lower limits that could be obtained given all possible combinations between the confidence intervals of each parameter of the model.

**Table S1. Parameter values estimated from experiments quantifying food competition (Experiment 1) or reproductive interference (Experiment 2).** Parameter values were estimated for each and across all five replicate populations. Values of $\lambda$ were computed as the mean daily production of daughters by females exposed to conspecific males only and grown in absence of food competition (one female per patch) in Experiments 1 and 2, then averaged between both experiments. Values of $\alpha$ and $\beta$ represent the negative effects of food competition and of reproductive interference, respectively, on offspring production per capita in Experiments 1 and 2, and were subsequently estimated using maximum likelihood. Confidence intervals (CI) are shown for the 95% level.

| | | **Replicate populations** | | | | | |
|---|---|---|---|---|---|---|---|
| **Parameters** | | **1** | **2** | **3** | **4** | **5** | **All** |
| $\lambda_{Tc}$ | Estimate | 5.73 | 5.14 | 6.81 | 5.07 | 5.11 | 5.57 |
| | Lower CI | 4.50 | 4.15 | 6.63 | 4.42 | 4.62 | 5.39 |
| | Upper CI | 6.96 | 6.12 | 6.99 | 5.72 | 5.61 | 5.75 |
| $\lambda_{Tu}$ | Estimate | 3.78 | 4.86 | 5.03 | 3.31 | 5.22 | 4.44 |
| | Lower CI | 1.66 | 2.98 | 3.92 | 1.46 | 4.40 | 2.88 |
| | Upper CI | 5.90 | 6.74 | 6.14 | 5.17 | 6.04 | 6.00 |
| $\alpha_{TcTc}$ | Estimate | 0.15 | 0.10 | 0.14 | 0.11 | 0.09 | 0.12 |
| | Lower CI | 0.10 | 0.06 | 0.11 | 0.08 | 0.07 | 0.10 |
| | Upper CI | 0.19 | 0.14 | 0.16 | 0.14 | 0.11 | 0.13 |
| $\alpha_{TcTu}$ | Estimate | 0.14 | 0.11 | 0.10 | 0.18 | 0.12 | 0.13 |
| | Lower CI | 0.07 | 0.06 | 0.05 | 0.11 | 0.07 | 0.10 |
| | Upper CI | 0.21 | 0.16 | 0.15 | 0.25 | 0.17 | 0.16 |
| $\alpha_{TuTu}$ | Estimate | 0.03 | 0.10 | 0.07 | 0.01 | 0.07 | 0.05 |
| | Lower CI | -0.01 | 0.04 | 0.03 | -0.01 | 0.03 | 0.04 |
| | Upper CI | 0.06 | 0.16 | 0.12 | 0.03 | 0.12 | 0.07 |
| $\alpha_{TuTc}$ | Estimate | 0.07 | 0.13 | 0.10 | 0.005 | 0.09 | 0.07 |
| | Lower CI | 0.03 | 0.05 | 0.05 | -0.02 | 0.04 | 0.05 |
| | Upper CI | 0.12 | 0.20 | 0.15 | 0.03 | 0.14 | 0.09 |
| $\beta_{TcTu}$ | Estimate | 1.23 | 0.03 | 0.03 | -0.13* | 0.22 | 0.21 |
| | Lower CI | 0.13 | -0.39 | -0.21 | -0.34 | -0.15 | -0.01 |
| | Upper CI | 2.33 | 0.45 | 0.27 | 0.09 | 0.58 | 0.43 |
| $\beta_{TuTc}$ | Estimate | 0.91 | 0.97 | 1.36 | -0.12* | 1.18 | 0.79 |
| | Lower CI | 0.27 | 0.15 | 0.49 | -0.42 | 0.04 | 0.45 |
| | Upper CI | 1.55 | 1.80 | 2.22 | 0.17 | 2.31 | 1.13 |

* As negative $\beta$ values are not supported by the current framework, population 4 was excluded from subsequent analyses involving this parameter.

**Table S2. Parameter values estimated from the experiment assessing the combined effects of food competition and reproductive interference (Experiment 3).** Parameter values were estimated for each and across all five replicate populations. Values for $\lambda$ were computed as the mean daily production of daughters by females exposed to conspecific males only and grown in absence of food competition (one female per patch). Values for $\alpha$ and $\beta$ represent the negative effects of food competition and of reproductive interference, respectively, on offspring production per capita, and were subsequently estimated using maximum likelihood, from data where females were exposed to heterospecific competitors for food and heterospecific mates. $\alpha$ parameters were estimated from data collected in the first generation (during which only food competition occurred), assuming no variation in the relationship between "intraspecific" and "interspecific" competition across experiments (*i.e.*, we used the relationships observed in Experiment 1; *e.g.*, $\alpha_{TcTc}$=0.83*$\alpha_{TcTu}$ and $\alpha_{TuTu}$=0.86*$\alpha_{TuTc}$ across all populations). $\beta$ parameters were subsequently estimated from data collected in the second and third generations of the experiment (nested approach). Confidence intervals (CI) are shown for the 95% level.

| | | Replicate populations | | | | | |
|---|---|---|---|---|---|---|---|
| **Parameters** | | **1** | **2** | **3** | **4** | **5** | **All** |
| $\lambda_{Tc}$ | Estimate | 1.47 | 1.8 | 3.00 | 2.90 | 2.73 | 2.38 |
| | Lower CI | 0.94 | 1.3 | 2.32 | 2.07 | 2 | 1.77 |
| | Upper CI | 2 | 2.3 | 3.66 | 3.73 | 3.47 | 2.99 |
| $\lambda_{Tu}$ | Estimate | 1.93 | 1.91 | 0.97 | 2.10 | 2.32 | 1.85 |
| | Lower CI | 1.26 | 1.2 | 0.42 | 1.25 | 1.46 | 1.39 |
| | Upper CI | 2.61 | 2.62 | 1.52 | 2.95 | 3.18 | 2.3 |
| $\alpha_{TcTc}$ | Estimate | 0.01 | 0.05 | 0.13 | 0.06 | 0.07 | 0.06 |
| | Lower CI | -0.01 | 0.04 | 0.10 | 0.02 | 0.04 | 0.05 |
| | Upper CI | 0.03 | 0.06 | 0.16 | 0.10 | 0.10 | 0.08 |
| $\alpha_{TcTu}$ | Estimate | 0.01 | 0.05 | 0.09 | 0.10 | 0.09 | 0.08 |
| | Lower CI | -0.01 | 0.04 | 0.07 | 0.03 | 0.05 | 0.06 |
| | Upper CI | 0.03 | 0.06 | 0.12 | 0.16 | 0.13 | 0.09 |
| $\alpha_{TuTu}$ | Estimate | 0.02 | 0.02 | -0.03* | 0.08 | 0.04 | 0.02 |
| | Lower CI | 0.01 | 0.003 | -0.04 | 0.06 | 0.01 | 0.01 |
| | Upper CI | 0.04 | 0.04 | -0.03 | 0.11 | 0.06 | 0.03 |
| $\alpha_{TuTc}$ | Estimate | 0.05 | 0.03 | -0.05* | 0.04 | 0.05 | 0.03 |
| | Lower CI | 0.02 | 0.005 | -0.06 | 0.03 | 0.01 | 0.02 |
| | Upper CI | 0.09 | 0.05 | -0.04 | 0.05 | 0.08 | 0.04 |
| $\beta_{TcTu}$ | Estimate | 0.10 | 0.20 | 0.39 | 0.49 | 0.35 | 0.30 |
| | Lower CI | -0.12 | 0.16 | 0.31 | 0.30 | 0.17 | 0.23 |
| | Upper CI | 0.31 | 0.25 | 0.48 | 0.68 | 0.53 | 0.37 |
| $\beta_{TuTc}$ | Estimate | 0.13 | 0.33 | 0.23 | 0.02 | 0.22 | 0.18 |
| | Lower CI | -0.08 | 0.16 | 0.05 | -0.14 | 0.01 | 0.10 |
| | Upper CI | 0.33 | 0.50 | 0.41 | 0.18 | 0.43 | 0.26 |

* As negative $\alpha$ values are not supported by the current framework, population 3 was excluded from subsequent analyses involving this parameter.

**Figure S1. Summary outcome of population dynamics in all simulated scenarios for all replicate populations.** The heat maps show the species predicted to remain after 20 generations, in simulations using different initial absolute densities and relative frequencies of both species. Rows of plots display the results obtained for each replicate population and across all replicates ("All data"), whereas columns of plots display which interaction(s) was(were) considered in the simulations: food competition only (FC), reproductive interference only (RI), using parameters estimated when both types of interaction occurred independently (Experiments 1 and 2; FC+RI), and when both occurred simultaneously (Experiment 3; FC*RI). Red: only Tc remains, Tu is excluded; Blue: only Tu remains, Tc is excluded.

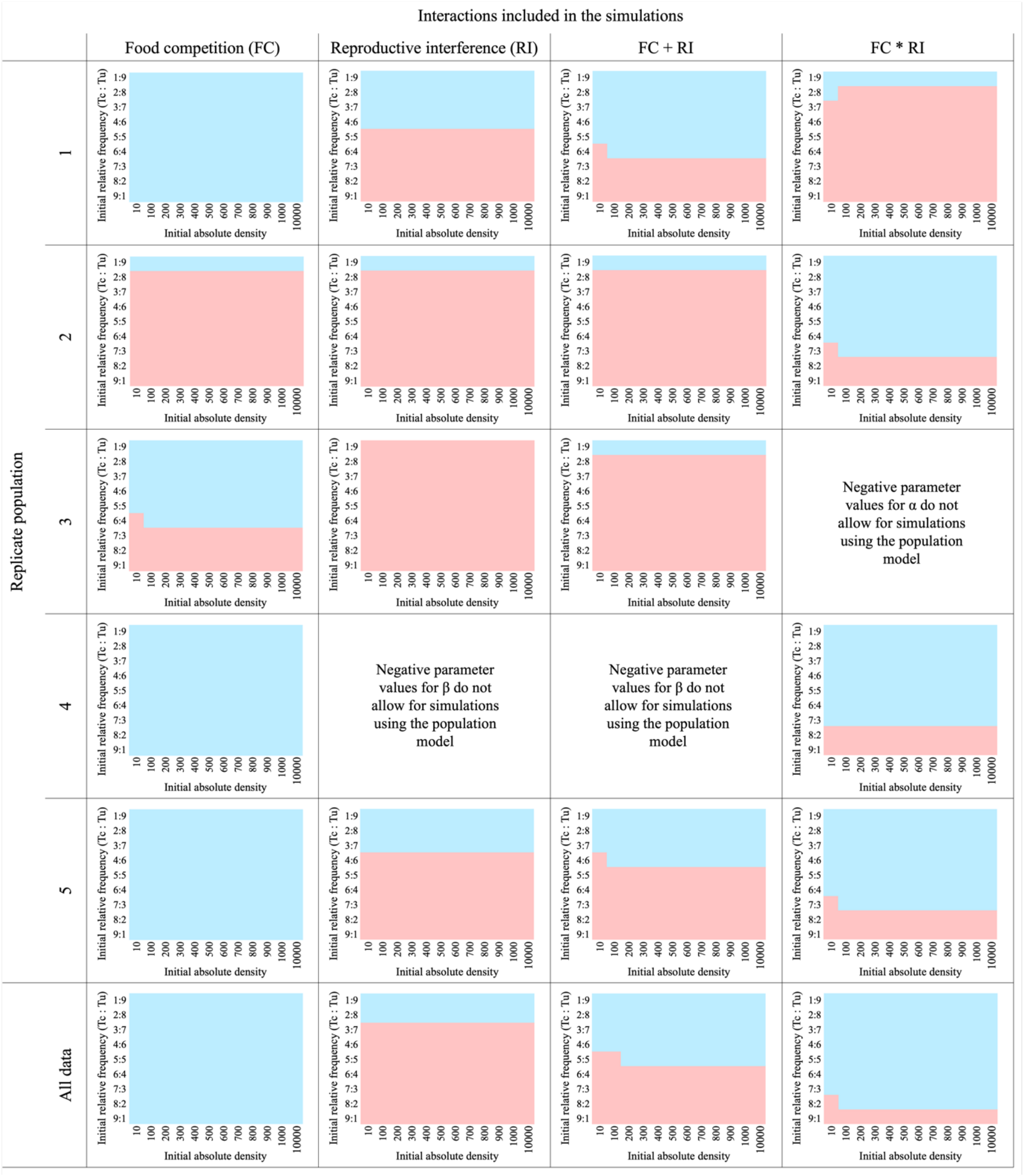

**Figure S2. Examples of simulated population dynamics assuming independent effects of food competition and reproductive interference.** Lines show changes in the number of females of each species across generations, when using parameter values estimated across all data in Experiment 1 and 2 ($\lambda_{Tu}$ = 4.44; $\lambda_{Tc}$ = 5.57; $\alpha_{TuTu}$ = 0.05; $\alpha_{TcTc}$ = 0.12; $\alpha_{TuTc}$ = 0.07; $\alpha_{TcTu}$ = 0.13; $\beta_{TuTc}$ = 0.79; $\beta_{TcTu}$ = 0.21). The outcome of simulations conducted for each replicate population is summarized in Figure S2. Rows of plots display increasing initial absolute densities of females from both species, whereas columns of plots display increasing initial frequency of Tc relative to Tu. Red lines: Tc; Blue lines: Tu.

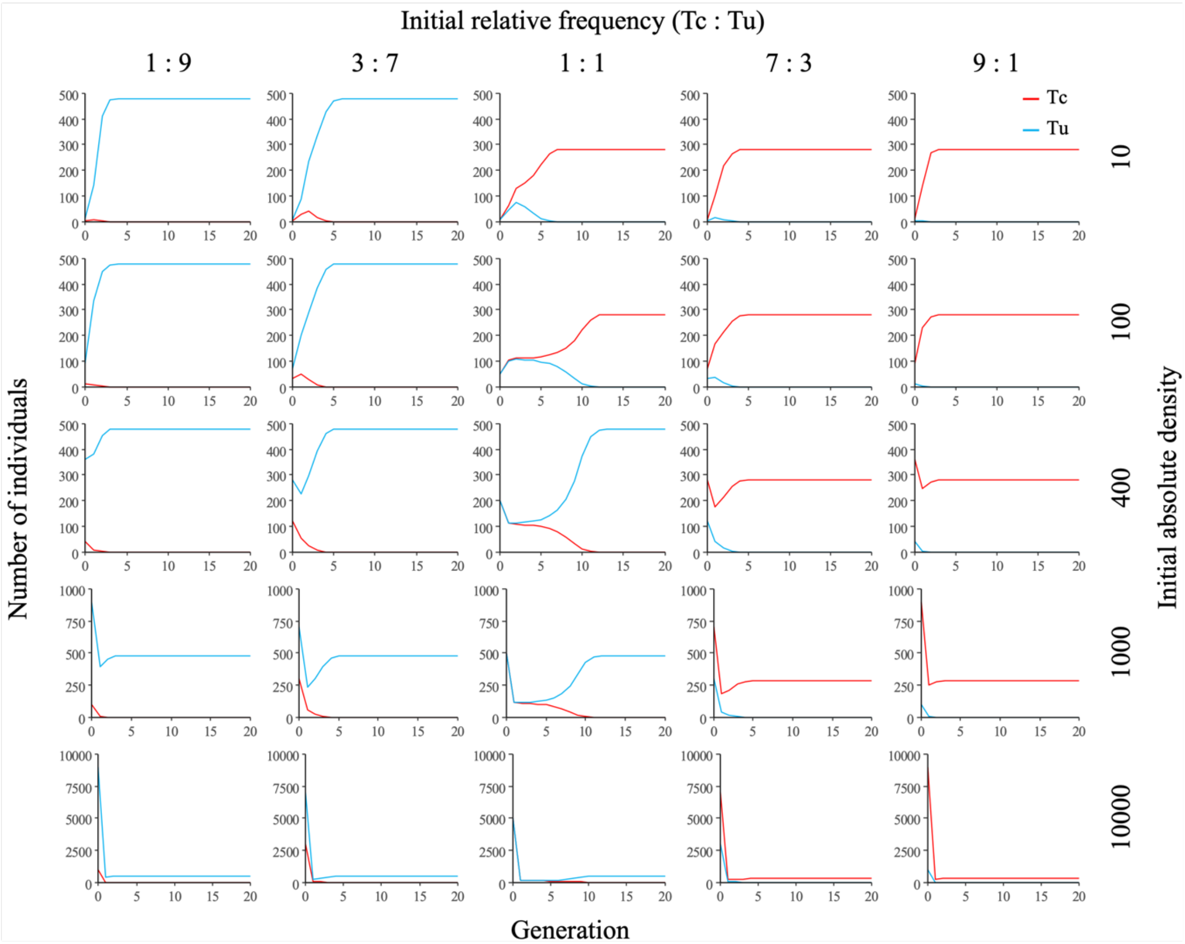

**Figure S3. Examples of simulated population dynamics allowing for interactive effects of food competition and reproductive interference.** Lines show changes in the number of females of each species across generations, when using parameter values estimated across all populations in Experiment 3 ($\lambda_{Tu}$ = 1.85; $\lambda_{Tc}$ = 2.38; $\alpha_{TuTu}$ = 0.02; $\alpha_{TcTc}$ = 0.06; $\alpha_{TuTc}$ = 0.03; $\alpha_{TcTu}$ = 0.08; $\beta_{TuTc}$ = 0.22; $\beta_{TcTu}$ = 0.32). The outcome of simulations conducted for each replicate population is summarized in Figure S2. Rows of plots display increasing absolute initial densities of females from both species, whereas columns of plots display increasing initial frequency of Tc relative to Tu. Red lines: Tc; Blue lines: Tu.

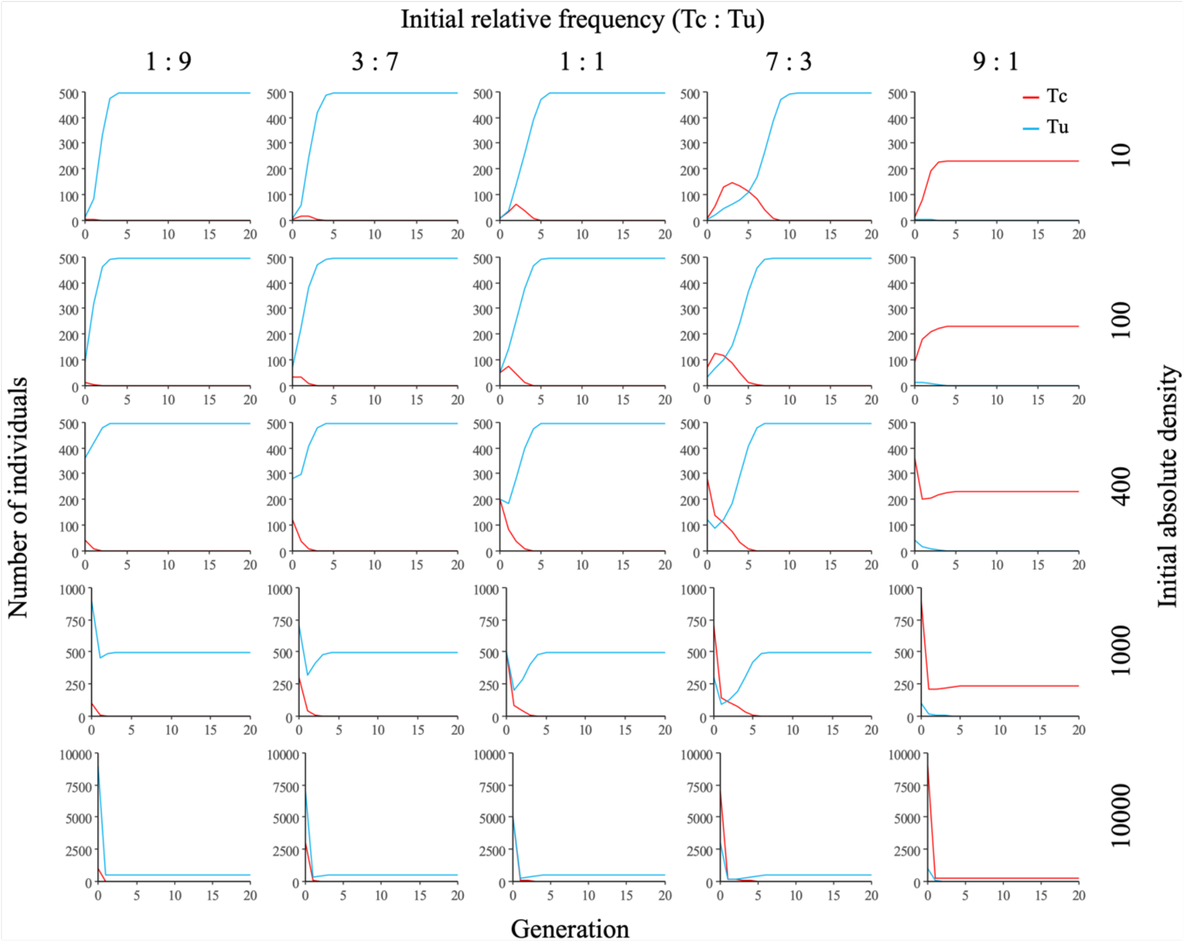

**Table S3. Estimated and predicted numbers and proportions of females in cage populations over 8 generations.** (A) Mean proportions (± confidence intervals) of Tc, Tu and hybrid female offspring ($P_{Tc}$, $P_{Tu}$ and $P_h$, respectively) estimated every two generations by phenotyping 50 randomly collected females in each experimental cage population (N=5), as well as the corresponding estimated numbers of sampled females ($S_{Tc}$, $S_{Tu}$ and $S_h$) used to start the next generation (with $S_{tot}$ = 400 females). (B) to (D) Predicted numbers ($N_{Tc}$, $N_{Tu}$ and $N_h$) and proportions ($P_{Tc}$, $P_{Tu}$ and $P_h$) of Tc, Tu, and hybrid female offspring, as well as predicted numbers of sampled females ($S_{Tc}$, $S_{Tu}$ and $S_h$) based on those proportions. Predictions were made using $\lambda$, $\alpha$, and $\beta$ parameter values obtained across all replicate populations in Experiments 1 and 2 (B), Experiment 3 (C), and values obtained from Experiment 3 but with $\alpha$/10 in Eq. S11 (used to compute $N_h$; D). Note that, in the experiment and all tested scenarios, hybrids were only formed from generation 2 onwards, as the females initially introduced in the cage populations were already mated with conspecific males.

| Generation | | 0 | 1 | 2 | 3 | 4 | 5 | 6 | 7 | 8 |
|---|---|---|---|---|---|---|---|---|---|---|
| **A) Estimated from experimental cage populations** | | | | | | | | | | |
| Mean ± CI | $P_{Tc}$ | 50 ± 0 | - | 10 ± 3 | - | 5 ± 4 | - | 0 ± 0 | - | 0 ± 0 |
| proportion | $P_{Tu}$ | 50 ± 0 | - | 51 ± 9 | - | 77 ± 8 | - | 94 ± 5 | - | 1 ± 0 |
| females (%) | $P_h$ | 0 ± 0 | - | 39 ± 8 | - | 18 ± 5 | - | 6 ± 5 | - | 0 ± 0 |
| Estimated | $S_{Tc}$ | 200 | - | 40 | - | 20 | - | 0 | - | 0 |
| No. sampled | $S_{Tu}$ | 200 | - | 204 | - | 308 | - | 376 | - | 400 |
| females | $S_h$ | 0 | - | 156 | - | 72 | - | 24 | - | 0 |
| **B) Predicted using parameter estimates from Experiments 1 and 2 (Independent effects of food competition and reproductive interference)** | | | | | | | | | | |
| Predicted | $N_{Tc}$ | - | 134 | 80 | 60 | 34 | 11 | <0.1 | <0.1 | <0.1 |
| No. offspring | $N_{Tu}$ | - | 202 | 163 | 200 | 269 | 363 | 444 | 472 | 475 |
| females | $N_h$ | - | 0 | 60 | 57 | 49 | 32 | 10 | 0.79 | <0.1 |
| Predicted | $P_{Tc}$ | 50 | 40 | 26 | 19 | 10 | 3 | < 1 | < 1 | < 1 |
| proportion | $P_{Tu}$ | 50 | 60 | 54 | 63 | 76 | 90 | 98 | 100 | 100 |
| females (%) | $P_h$ | 0 | 0 | 20 | 18 | 14 | 8 | 2 | < 1 | < 1 |
| Predicted | $S_{Tc}$ | 200 | 134 | 80 | 60 | 34 | 11 | 1 | 0 | 0 |
| No. sampled | $S_{Tu}$ | 200 | 202 | 163 | 200 | 269 | 358 | 390 | 399 | 400 |
| females | $S_h$ | 0 | 0 | 60 | 57 | 49 | 31 | 9 | 1 | 0 |
| **C) Predicted using parameter estimates from Experiment 3 (Interactive effects of food competition and reproductive interference)** | | | | | | | | | | |
| Predicted | $N_{Tc}$ | - | 98 | 37 | 8 | 0.24 | <0.1 | <0.1 | <0.1 | <0.1 |
| No. offspring | $N_{Tu}$ | - | 207 | 258 | 359 | 439 | 446 | 446 | 446 | 446 |
| females | $N_h$ | - | 0 | 30 | 17 | 4 | 0.1 | <0.1 | <0.1 | <0.1 |
| Predicted | $P_{Tc}$ | 50 | 32 | 11 | 2 | < 1 | < 1 | < 1 | < 1 | < 1 |
| proportion | $P_{Tu}$ | 50 | 68 | 79 | 94 | 99 | 100 | 100 | 100 | 100 |
| females (%) | $P_h$ | 0 | 10 | 5 | 1 | < 1 | < 1 | < 1 | < 1 | < 1 |
| Predicted | $S_{Tc}$ | 200 | 98 | 37 | 8 | 0 | 0 | 0 | 0 | 0 |
| No. sampled | $S_{Tu}$ | 200 | 207 | 258 | 359 | 397 | 400 | 400 | 400 | 400 |
| females | $S_h$ | 0 | 0 | 30 | 17 | 3 | 0 | 0 | 0 | 0 |
| **D) Predicted using parameter estimates from Experiment 3, but hybrids 10 times less sensitive to food competition (Interactive effects of food competition and reproductive interference)** | | | | | | | | | | |
| Predicted | $N_{Tc}$ | - | 98 | 37 | 7 | 0.25 | <0.1 | <0.1 | <0.1 | <0.1 |
| No. offspring | $N_{Tu}$ | - | 207 | 258 | 350 | 450 | 442 | 445 | 446 | 446 |
| females | $N_h$ | - | 0 | 189 | 105 | 25 | 0.8 | <0.1 | <0.1 | <0.1 |
| Predicted | $P_{Tc}$ | 50 | 32 | 8 | 2 | < 1 | < 1 | < 1 | < 1 | < 1 |
| proportion | $P_{Tu}$ | 50 | 68 | 53 | 76 | 94 | 100 | 100 | 100 | 100 |
| females (%) | $P_h$ | 0 | 0 | 39 | 23 | 6 | < 1 | < 1 | < 1 | < 1 |
| Predicted | $S_{Tc}$ | 200 | 98 | 31 | 6 | 0 | 0 | 0 | 0 | 0 |
| No. sampled | $S_{Tu}$ | 200 | 207 | 213 | 303 | 378 | 399 | 400 | 400 | 400 |
| females | $S_h$ | 0 | 0 | 156 | 91 | 22 | 1 | 0 | 0 | 0 |

**Figure S4. Predicted dynamics of *Tetranychus urticae* (Tu), *T. cinnabarinus* (Tc), and of their hybrids, in cage populations over 8 generations.** Predicted proportions of females assuming that hybrid females are as strongly affected by food competition as purebred females, with coefficients measured either when food competition and reproductive interference acted separately (A), or simultaneously (B; parameters estimated across all replicate populations, from Experiments 1 and 2, or from Experiment 3, respectively). Coloured lines and shaded areas show predicted proportions and associated upper and lower limits that could be obtained given all possible combinations between the 95% confidence intervals of each parameter of the model. Note that all confidence intervals fully overlap from generation 5 onward in panel A. Blue: Tu females; Red: Tc females; Purple: hybrid females.

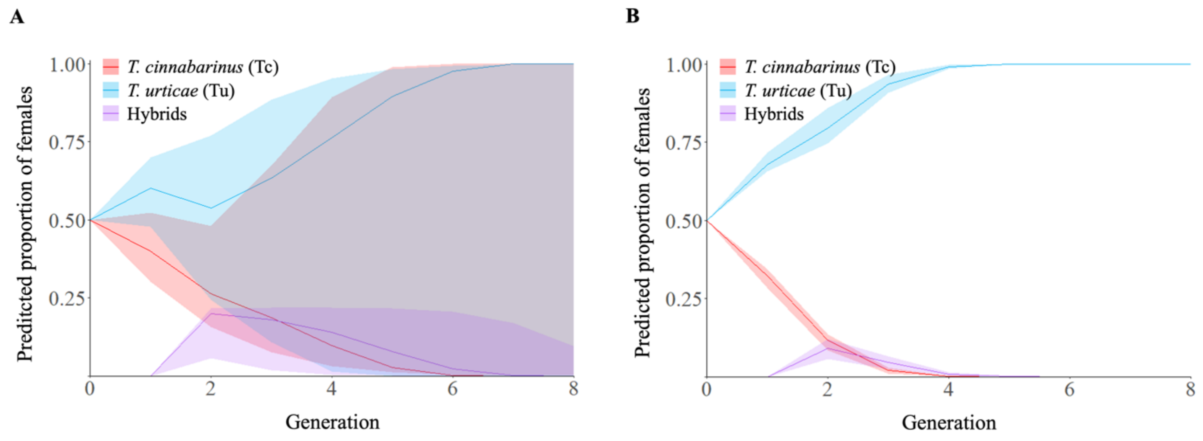

**Table S4. Coefficients obtained for each linear regression between observed cage population dynamics and predictions based on different scale factors of the alpha parameters for hybrid sensitivity to food competition.** The table shows R-squared, intercept, regression coefficient, and Akaike information criterion (AIC) values obtained for each linear regression performed between the observed proportions of females and those predicted by the model, for a range of scale factors of the $\alpha$ parameters in Eq. S11. Shaded cells indicate the scale factor beyond which the combination of R-squared, intercept and regression coefficient no longer improved (and the intercept is already very close to its minimum value).

| Scale factor of $\alpha$ | Intercept | Regression coefficient | R-Squared | AIC |
|---|---|---|---|---|
| 1 | 0.064 | 0.81 | 0.89 | 32.69 |
| 2 | 0.055 | 0.83 | 0.92 | 32.78 |
| 3 | 0.049 | 0.85 | 0.93 | 32.87 |
| 4 | 0.045 | 0.86 | 0.94 | 32.94 |
| 5 | 0.042 | 0.87 | 0.95 | 33.01 |
| 6 | 0.040 | 0.88 | 0.96 | 33.08 |
| 7 | 0.038 | 0.89 | 0.96 | 33.14 |
| 8 | 0.036 | 0.89 | 0.96 | 33.19 |
| 9 | 0.035 | 0.89 | 0.96 | 33.25 |
| 10 | 0.035 | 0.90 | 0.96 | 33.29 |
| 11 | 0.034 | 0.90 | 0.97 | 33.34 |
| 12 | 0.033 | 0.90 | 0.97 | 33.38 |
| 13 | 0.033 | 0.90 | 0.97 | 33.41 |
| 14 | 0.033 | 0.90 | 0.97 | 35.24 |
| 15 | 0.032 | 0.90 | 0.97 | 35.24 |
| 16 | 0.032 | 0.90 | 0.96 | 35.24 |
| 17 | 0.032 | 0.90 | 0.96 | 35.25 |
| 18 | 0.032 | 0.90 | 0.96 | 35.25 |
| 19 | 0.032 | 0.90 | 0.96 | 35.26 |
| 20 | 0.032 | 0.90 | 0.96 | 35.26 |
| 21 | 0.032 | 0.90 | 0.96 | 35.27 |
| 22 | 0.032 | 0.91 | 0.96 | 35.27 |
| 23 | 0.032 | 0.91 | 0.96 | 35.27 |
| 24 | 0.032 | 0.91 | 0.96 | 43.61 |
| 25 | 0.032 | 0.91 | 0.96 | 43.60 |
| 26 | 0.032 | 0.91 | 0.96 | 43.59 |
| 27 | 0.032 | 0.91 | 0.96 | 43.59 |
| 28 | 0.032 | 0.91 | 0.96 | 43.58 |
| 29 | 0.032 | 0.91 | 0.96 | 43.57 |
| 30 | 0.032 | 0.91 | 0.96 | 43.57 |
